# IRE1α undergoes a dynamic filament assembly cycle governed by unfolded proteins and BiP

**DOI:** 10.64898/2026.10.19.760176

**Authors:** Melissa Méndez-Moran, Maxence Le-Vasseur, Daniel N. Itzhak, Wiktoria Sadowska, G. Elif Karagöz, Peter Walter, David A. Agard

**Affiliations:** University of California San Francisco, Biochemistry and Biophysics, San Francisco, CA; Altos Labs Inc, Institute of Science, Redwood City, CA; Max Perutz Labs and Medical University of Vienna, Vienna, Austria; current address: Buck Institute for Research on Aging, Novato, CA; Biohub, Redwood City, CA

## Abstract

The endoplasmic reticulum is a major hub of protein synthesis, modification and trafficking that relies on the Unfolded Protein Response (UPR) to balance its folding capacity. IRE1, the most conserved of the UPR sensors, undergoes a change in oligomeric state upon activation that propagates the stress signal from the ER lumen to its cytoplasmic kinase/RNase domains. Here we show that opposing inputs from unfolded proteins and the ER chaperone machinery govern a dynamic cycle of IRE1 assembly. A *bona fide* unfolded ER client directly binds the IRE1 lumenal domain (LD) and promotes higher-order oligomerization. Model peptides further reveal that full assembly into LD filaments depends on ligand length, sequence, and IRE1’s oligomerization interfaces. The ER Hsp70, BiP, counterbalances this process through two distinct mechanisms: i) an ATP-independent interaction via BiP’s nucleotide-binding domain buffers early IRE1 assembly by engaging its oligomerization interface, whereas ii) ATP-regulated BiP/ERDJ machinery actively dismantles assembled filaments. Together, our findings reconcile direct unfolded protein recognition with BiP-mediated regulation to establish LD oligomerization as a point of integration through which unfolded protein load and chaperone availability dynamically regulate IRE1 assembly state and signalling.

## Introduction

Cells invest substantial resources into keeping their proteomes folded and functional. The Unfolded Protein Response (UPR) responds to perturbations in the protein folding capacity of the endoplasmic reticulum (ER) by upregulating chaperone-assisted protein folding, ER-associated degradation of misfolded proteins (ERAD), and downregulating protein translation (1, 2). However, if the ER stress cannot be resolved, unmitigated UPR signalling initiates pro-apoptotic pathways that eliminate maladaptive cells (3-6). By adjusting its signalling output to the severity and persistence of ER stress, the UPR makes life/death decisions for the cell that place it at the centre of diverse human pathologies including neurodegeneration (7-9), cancer (10-12), and metabolic disease (13, 14).

The most conserved branch of the UPR is driven by the transmembrane kinase/RNase sensor IRE1-alpha (15, 16). Activation of IRE1 is coupled to the oligomerization of its ER lumenal domain (LD), which drives proper alignment of its cytoplasmic domains, enabling trans-autophosphorylation and endonuclease activity (17-20). Active IRE1 then splices mRNA encoding the transcription factor XBP1, facilitating upregulation of UPR target genes including ER chaperones and components of ERAD machinery (21, 22). Additionally, active IRE1 can cleave ER-localised mRNAs through regulated IRE1-dependent decay (RIDD/RIDDLE) to further alleviate ER folding load (23, 24). Assembly of the core folded segment of the IRE1 lumenal domain resolved in the human crystal structure (cLD: aa24-389) is mediated through two distinct interfaces: IF1 stabilizes head-to-head dimer formation and IF2 mediates tail-to-tail association into higher-order oligomers (25, 26) (Fig. 2A). Both are critical for stress-induced IRE1 clustering and their mutation results in dysregulated signalling (20, 26).

A central question in IRE1 regulation remains how its cLD detects changes in ER protein-folding status and how this in turn promotes oligomerization. Early studies identified an inverse correlation between IRE1 activity and association with the ER molecular chaperone BiP (27, 28), leading to a model in which IRE1 indirectly senses ER stress by monitoring chaperone availability through an ATP-driven client interaction with the BiP substrate-binding domain (SBD). In this model, the co-chaperone ERDJ4 assists BiP in sequestering inactive IRE1 monomers until accumulating unfolded proteins (UPs) “titrate” BiP away, allowing for IRE1-cLD assembly (29, 30). However, IRE1 mutants defective in BiP association display dysregulated, but not constitutive signalling (31, 32), suggesting that BiP dissociation alone does not fully account for IRE1 activation. By contrast, UPs have been proposed to act directly as activating ligands by binding the IRE1-cLD (26, 33, 34). This notion emerged from observations of structural homology between the IRE1-cLD dimer interface and the major histocompatibility complex (MHC) peptide-binding groove and *in vitro* data showing that purified IRE1-cLD binds unfolded polypeptides (26). More recently, a non-canonical mode of BiP engagement with IRE1 was identified when an ATP-independent and nucleotide-binding domain (NBD)-driven interaction with IRE1-cLD was measured *in vitro* (35). Engagement of unfolded proteins by the BiP SBD was proposed to allosterically release the NBD from IRE1 during ER stress (36, 37), but how this NBD-driven interaction affects IRE1-cLD’s propensity for self-assembly or intersects with direct UP binding remains unknown. IRE1 additionally integrates stress inputs directly from its lipid membrane environment (38, 39) and protein folding load via interactions with the ER-docked ribosome and/or SEC61 translocon (40-43), emphasizing the complexity of its surveillance over the ER’s protein status.

Together, these observations suggest a multi-layered regulatory network for IRE1 and the broader UPR, necessitating investigation of how mechanisms historically considered as alternatives interact to yield a tuneable and sensitive stress response. Here we reconstitute key players in IRE1 regulation to dissect how unfolded proteins and distinct modes of BiP engagement govern IRE1-cLD oligomerization. We establish a mechanistic framework for understanding how IRE1 assembly is dynamically promoted, buffered, and resolved.

## Results

### A *bona fide* unfolded ER client protein induces IRE1-cLD filaments

While a growing body of evidence supports the ability of IRE1 to bind unfolded proteins directly, a limited number of physiologically plausible clients have been identified. To determine whether direct binding of an unfolded protein client is coupled to changes in IRE1 assembly state, we asked whether unfolded bovine pancreatic trypsin inhibitor (BPTI), a 58-amino-acid *bona fide* secretory protein that folds compactly in the ER lumen, would influence its oligomeric state. BPTI showed two regions of binding in its primary sequence (Fig. 1A, SuppFig. 1A) on peptide arrays probed with IRE1-cLD. We designed a fluorescently labelled 48-amino-acid peptide (BPTI-ΔC) spanning both regions, in which we substituted the remaining 4 cysteines with alanines. Since disulfide bonds confer the majority of PTI’s folding energy, this substitution results in an unfolded protein, as previously reported (44, 45). BPTI-ΔC bound to IRE1-cLD as assessed by fluorescence anisotropy (K_1/2_ = 18.5 µM) (Fig. 1B), with an affinity comparable to previously reported unfolded protein model substrates (26, 46). BPTI-ΔC binding to IRE1-cLD was not outcompeted by disulfide-bonded, unlabeled BPTI, indicating that the binding of BPTI-ΔC results from its unfoldedness (Fig. 1C).

**Figure 1.**
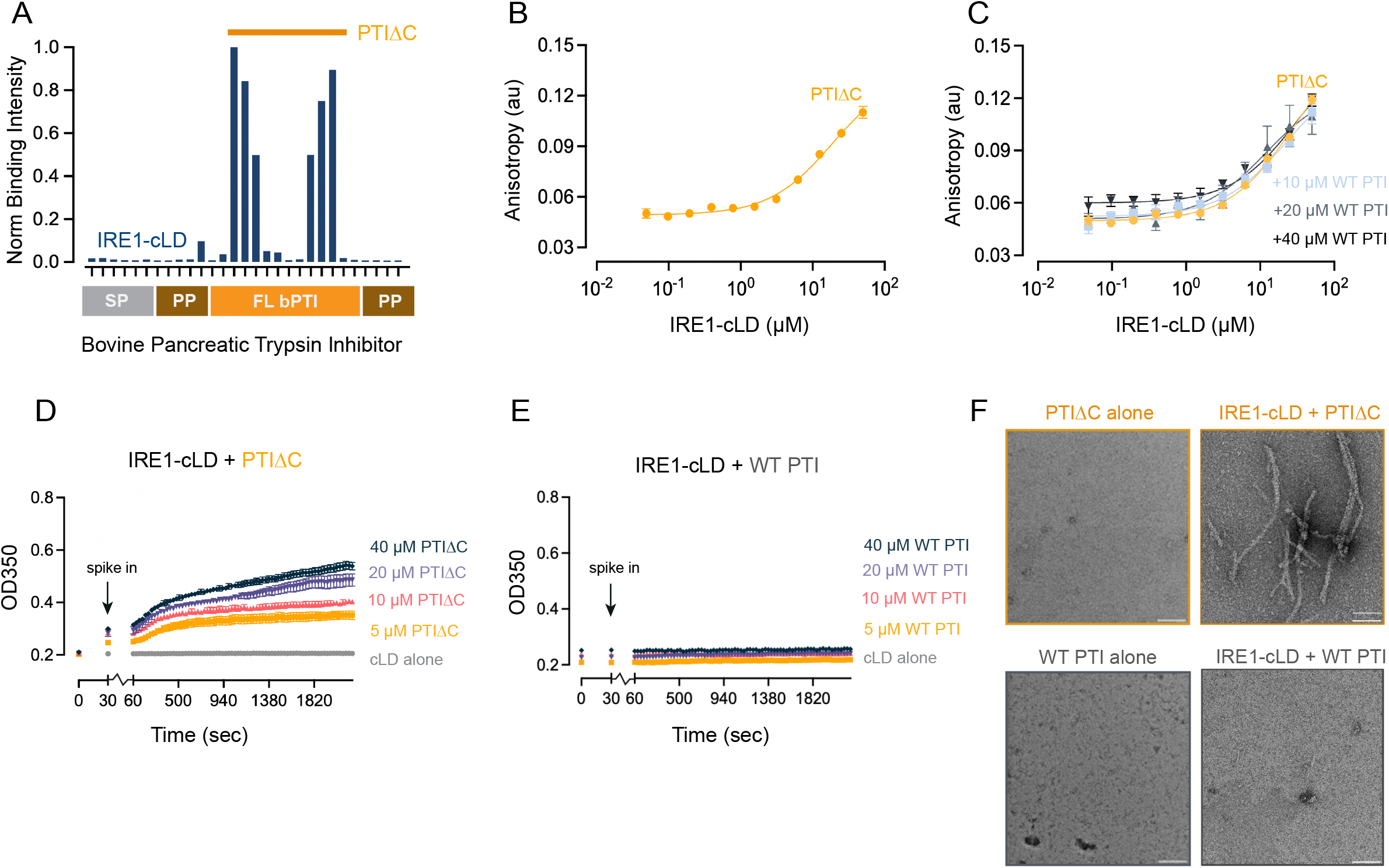
Unfolded bovine PTI induces assembly of IRE1-cLD into filaments. A. Binding intensities of IRE1-cLD to 18-mer peptides tiled along the sequence of bovine pancreatic trypsin inhibitor. The binding regions used to generate PTIΔC are indicated. The topology of full length PTI is shown below the horizontal axis (SP: signal peptide, PP: pro-peptide). B. IRE1-cLD binds to PTIΔC with a K_1/2_ ≈ 18.5 µM, as determined by fluorescence anisotropy (sequence: LEPPYTGPAKARIIRYFYNAKAGLAQTFVYGGARAKRNNFKSAE). C. Increasing concentrations of WT folded PTI do not impair binding of IRE1-cLD to PTIΔC as determined by fluorescence anisotropy. D. Light scattering of 5 µM IRE1-cLD in the presence of increasing concentrations of the unfolded PTIΔC peptide (0-40 µM) over a 1 hr period at 25°C. E. Light scattering at UV 350 nm of 5 µM WT IRE1 cLD in the presence of increasing concentrations of full length folded pancreatic trypsin inhibitor (0-40 µM) over a 1 hr period at 25°C. F. Negative stain TEM micrographs (42kx mag and -1.04 µm defocus) of WT IRE1 cLD in the presence of either FL PTI or PTIΔC. Filaments are only observed when PTI is unfolded.

**Figure 2.**
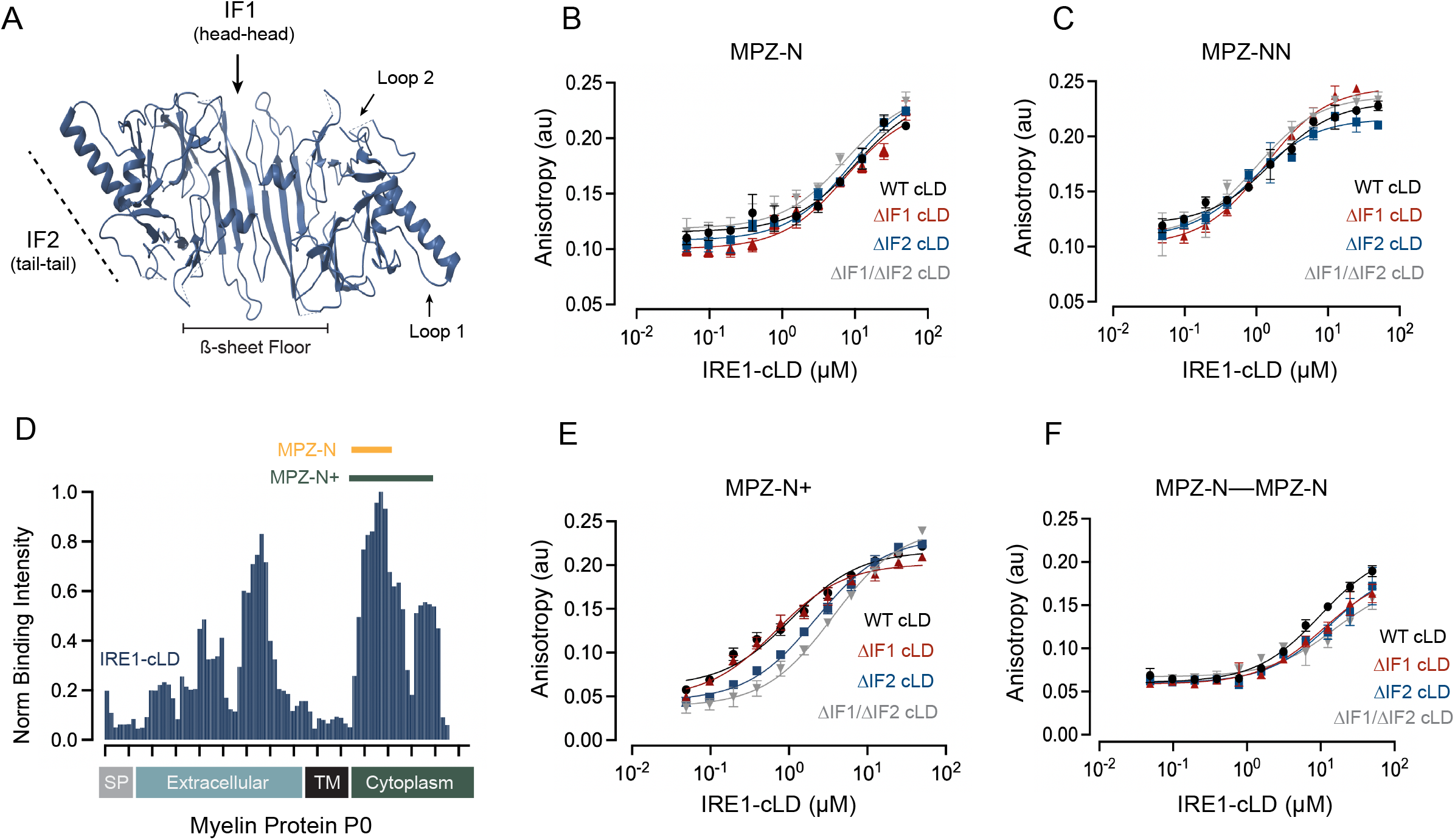
MPZ-derived peptides bind IRE1-cLD in a multivalent and IF2-dependent manner. A. Notable regions of human IRE1-cLD are depicted on the dimer crystal structure from PDB deposition 2HZ6 (25). The beta-sheet floor constituting the base of the “MHC-like groove” (highlighted in black) bridges across two cLD dimers at IF1. B. IRE1-cLD binding to the unfolded peptide MPZ-N (sequence: LIRYAWLRRQAA) is not impaired by cLD mutants compromising their ability to form IF1 and/or IF2 as determined by fluorescence anisotropy (Table of K_1/2_ in SuppFig. 2). C. IRE1-cLD interface mutants do not diminish its affinity for the tandem binding site peptide MPZ-NN (sequence: LIRYAWLRRQAAGSLIRYAWLRRQAA) as shown by fluorescence anisotropy (Table of K_1/2_ values in SuppFig. 2). D. Binding intensities of IRE1-cLD to 18-mer peptides tiled along the full native sequence of MPZ adapted from (26). The topology of MPZ is shown below the horizontal axis (SP: signal peptide, TM: transmembrane domain). The cytoplasmic IRE1 binding region identified by the authors displays a bimodal peak extending beyond the MPZ-N peptide sequence. Intensities corresponding to the MPZ-N and MPZ-N+ binding regions are indicated. E. IRE1-cLD binding to the extended unfolded peptide MPZ-N+ (sequence: LIRYAWLRRQAALQRRISAMEGSKGKLHKPGKDASKRGRQTPVLYAML) shows a defect when incubated with cLD mutants incapable of forming IF2 (Table of K_1/2_ values in SuppFig. 2). F. IRE1-cLD variant binding to the unfolded MPZ-N––MPZ-N peptide (sequence: LIRYAWLRRQAAGSGSGSGSGSGSGSGSGSGSGSLIRYAWLRRQAA) shows a reduced affinity and negligible defects in the presence of IRE1-cLD interface mutants (Table of K_1/2_ values in SuppFig. 2).

In the absence of BPTI-ΔC, IRE1-cLD exists as a homogeneous dimer at 5 µM as observed via native mass spectrometry (MS) analysis, which at higher concentrations assemble into tetramers (SuppFig. 1B, C). To determine whether IRE1-cLD’s oligomerization state is affected by BPTI-ΔC binding, we measured turbidity by light scattering at 350 nm in the presence of increasing BTPI-ΔC concentrations (Fig. 1D). We observed a substantial increase in light scattering induced by BPTI-ΔC over a 30 min time course, whereas disulfide-bonded, folded BPTI had no effect (Fig. 1E). Remarkably, inspection of the mixtures by negative stain electron microscopy (EM) revealed that BTPI-ΔC, but not folded BTPI, induced assembly of IRE1-cLD into filaments (Fig. 1F). Taken together, our findings show that binding to unfolded proteins drives IRE1-cLD’s oligomerization state.

### Unfolded peptide length and sequence together determine engagement with IRE1-cLD

Having established that an unfolded ER client can promote IRE1-cLD oligomerization, we sought to define how unfolded ligands engage the IRE1-cLD and whether binding depends on its known self-association interfaces. To this end, we employed fluorescently labelled peptides derived from Myelin Protein Zero (MPZ), which was established as a powerful proxy for UP binding to the IRE1-cLD (26). Unlike BPTI-ΔC, MPZ contains a previously characterized IRE1-cLD binding sequence with known sequence preferences. Although this sequence is not present in the ER lumen under physiological conditions, its prior characterization together with its constitutive lack of structure (47) and more tractable affinity for IRE1-cLD *in vitro* makes MPZ a valuable model for mechanistic dissection of UP binding. Consistent with previous work, a 12-amino-acid peptide (“MPZ-N”) bound to purified human wildtype IRE1-cLD as measured by fluorescence polarization anisotropy (WT; K_1/2_ = 10.39 µM) (Fig. 2B, WT). To assess the extent of the MPZ-N binding site, we performed the binding assays with mutant cLDs in which we disrupted the interfaces known to mediate IRE1-cLD head-to-head dimerization (cLD-ΔIF1), tail-to-tail oligomerization (cLD-ΔIF2), or both (cLD-ΔIF1/2). WT cLD purified as a dimer (SuppFig. 2A). As expected, cLD-ΔIF1 and cLD-ΔIF1/2 purified as monomers, whereas cLD-ΔIF2 remained a dimer (SuppFig. 2A). Unexpectedly, all three mutants bound MPZ-N with indistinguishable affinity (Fig. 2B), indicating that the measured IRE1-cLD•peptide interaction is not compromised by disruption of either interface. Hence, the binding site is likely contained within a single IRE1-cLD monomer. We obtained the same results for a 26 amino acid tandem repeat of MPZ-N (MPZ-NN) (Fig. 2C). As previously noted (23), MPZ-NN binds ∼10-fold more tightly (K_1/2_ = 1.75 µM) than MPZ-N, suggesting either a more extended interface or statistically favored rebinding enabled by the repeat structure of the peptide. In either case, the binding site is still fully contained in an IRE1-cLD monomer.

Extending the peptide size to match the length of BPTI-ΔC, we generated a 48-amino-acid peptide that includes further sequence elements from MPZ. This sequence was selected based on a previously reported MPZ tiling blot probed with IRE1-cLD (Fig. 2D) which, like BTPI-ΔC, shows an extended region of IRE1-binding (MPZ-N+). MPZ-N+ displayed a further increase in affinity for IRE1-cLD (K_1/2_ = 1.08 µM) (Fig. 2E). Scrambling its sequence abolished binding to IRE1-cLD (SuppFig. 2B), indicating that the interaction depends on sequence-specific elements. In contrast to MPZ-N and MPZ-NN, MPZ-N+ binding to cLD interface mutants revealed a defect in mutants that cannot engage in tail-to-tail oligomerization (cLD-ΔIF2 and cLD-ΔIF1/2) but showed no impairment for those that cannot dimerize head-to-head (ΔIF1) (Fig. 2E). These observations suggest that either (i) the longer MPZ-N+ peptide bridges across the IF2 oligomerization interface to interact with two IRE1-cLD monomers, or (ii) that binding of MPZ-N+ remains contained within one monomer and that binding is sensitive to IF2 mutation due to secondary MPZ-N+ contacts localized to IF2.

To begin to distinguish between these possibilities, we next assessed the importance of the full peptide length. To this end, we constructed a peptide composed of two MPZ-N modules connected with a 22aa glycine-serine linker (MPZ-N––MPZ-N), which renders the peptide the same overall length as MPZ-N+. We found that MPZ-N––MPZ-N bound IRE1-cLD with an affinity similar to that of MPZ-N (K_1/2_ = 11.58 µM vs. K_1/2_ = 10.39 µM, respectively), suggesting that the glycine-serine linker renders the second MPZ-N motif irrelevant. Furthermore, akin to MPZ-N, MPZ-N––MPZ-N showed only marginal defects in binding to cLD IF2 mutants (Fig. 2F). Taken together, these data are consistent with the notion that the additional sequence elements appended in MPZ-N+ directly adjacent to the short MPZ-N sequence lead to its increased binding affinity and IF2-dependence, possibly through auxiliary contacts at IF2 that can be accommodated within a single IRE1-cLD monomer.

### MPZ-N+ facilitates the assembly of IRE1-cLD into filaments

Given the contribution of IF2 to MPZ-N+ binding affinity and the notion that IF2 links IRE1-cLD dimers in formation of larger oligomers (25, 26), we next asked whether the distinct binding properties of MPZ peptides translate into differences in their capacity to promote IRE1-cLD assembly. Akin to BPTI-ΔC, MPZ-N+ induced IRE1-cLD oligomerization in our light scattering assay (Fig. 3A). Similarly, negative stain EM revealed that MPZ-N+ induced the formation of IRE1-cLD filaments of similar dimensions as those induced by BPTI-ΔC (Fig. 3E). The filaments measured 90 Å in diameter and, at higher magnification, showed a substructure of repetitively arranged modules spaced at approximately 167 Å. Both light scattering and filament assembly were dependent on IF1 and IF2 (Fig. 3B, C, E).

**Figure 3.**
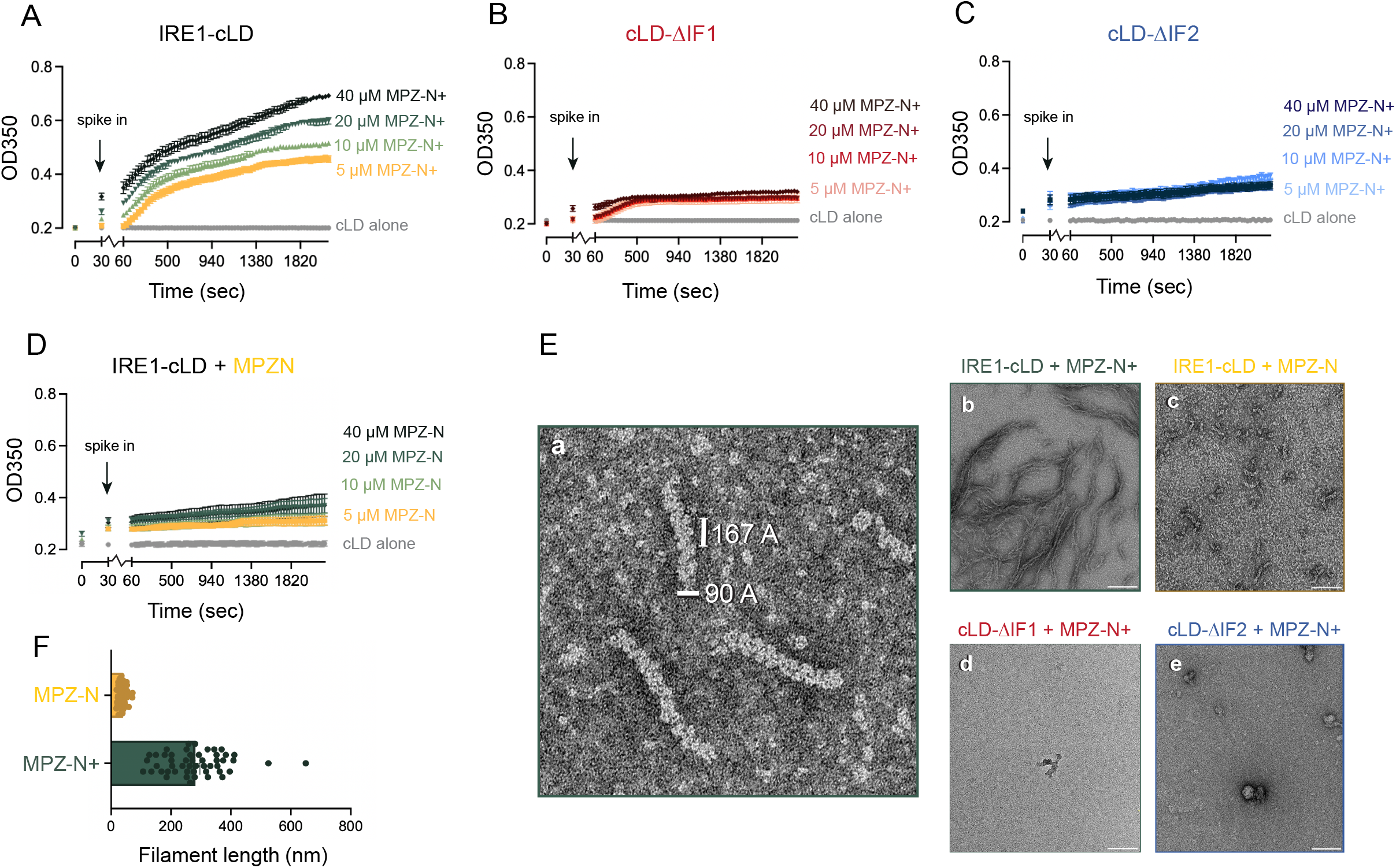
The extended MPZ-N+ sequence also induces IRE1-cLD filament formation. A. Light scattering at UV 350 nm of 5 µM IRE1-cLD in the presence of increasing concentrations of MPZ-N+ (0-40 µM) over a 1 hr period at 25°C. B. Light scattering of 5 µM of the dimerization-deficient IRE1-cLD mutant (ΔIF1) in the presence of increasing concentrations of MPZ-N+ (0-40 µM) over a 1 hr period at 25°C. C. Light scattering of 5 µM of the oligomerization-deficient IRE1-cLD mutant (ΔIF2) in the presence of increasing concentrations of MPZ-N+ (0-40 µM) over a 1 hr period at 25 °C. D. Light scattering of IRE1-cLD in the presence of increasing concentrations of the short peptide MPZ-N (0-40 µM) over a 1 hr period at 25°C. E. Negative stain TEM micrographs (a: 100kx mag/-1.04 µm defocus; b-e: 42kx mag and -1.04 µm defocus) of 5 µM IRE1-cLD’s oligomerization status induced in the presence or absence of 20 µM unfolded MPZ peptides. Significant cLD filament assembly is observed only when WT IRE1-cLD is incubated with the extended domain sequence MPZ-N+. Ordered repeating units are visible, and a diameter of 90 A is indicated. F. Bar chart of IRE1-cLD filament lengths measured in the presence of either MPZ-N or MPZ-N+ peptide. Measurements of 50 filaments from 10 (MPZ-N) or 14 (MPZ-N+) micrographs were performed using IMOD’s 3dmod program (79).

By contrast to MPZ-N+, the scrambled version of the peptide (the non-binder control; SuppFig. 2B) did not induce light scattering (SuppFig. 3D), indicating that light scattering resulted from specific IRE1-cLD ligand-binding and was not due to molecular crowding or other nonspecific interactions. Furthermore, the shorter MPZ-N peptide also failed to induce light scattering, confirming that the additional sequence elements in MPZ-N+ are required (Fig. 3D). However, negative stain EM revealed that MPZ-N did provoke formation of small IRE1-cLD oligomers (Fig. 3E,F). Similarly, the MPZ-N––MPZ-N peptide showed only a modest increase in light scattering (SuppFig. 3E) and, like MPZ-N, the formation of shorter oligomers (SuppFig. 3F). Taken together, our findings demonstrate that the extended interaction with IRE1 monomers enabled by additional sequence elements contained in MPZ-N+ plays a major role in promoting an oligomerization-favored state in IRE1-cLD.

### The BiP NBD buffers early-stage IRE1-cLD oligomerization to prevent filament assembly

Ample evidence supports BiP as a second major regulator in IRE1 stress sensing (27, 29, 32). To investigate the interplay between UP and BiP binding, we next asked whether the purified nucleotide-binding domain of BiP (BiP-NBD) competes or synergizes with either IRE1-cLD peptide binding or oligomerization. We saw no effect on MPZ-N binding in the presence of BiP-NBD bound to either ADP or the non-hydrolyzable analogue AMPPNP (Fig. 4A), indicating that the two components bind IRE1-cLD independently regardless of the nucleotide-bound state of the BiP-NBD.

**Figure 4.**
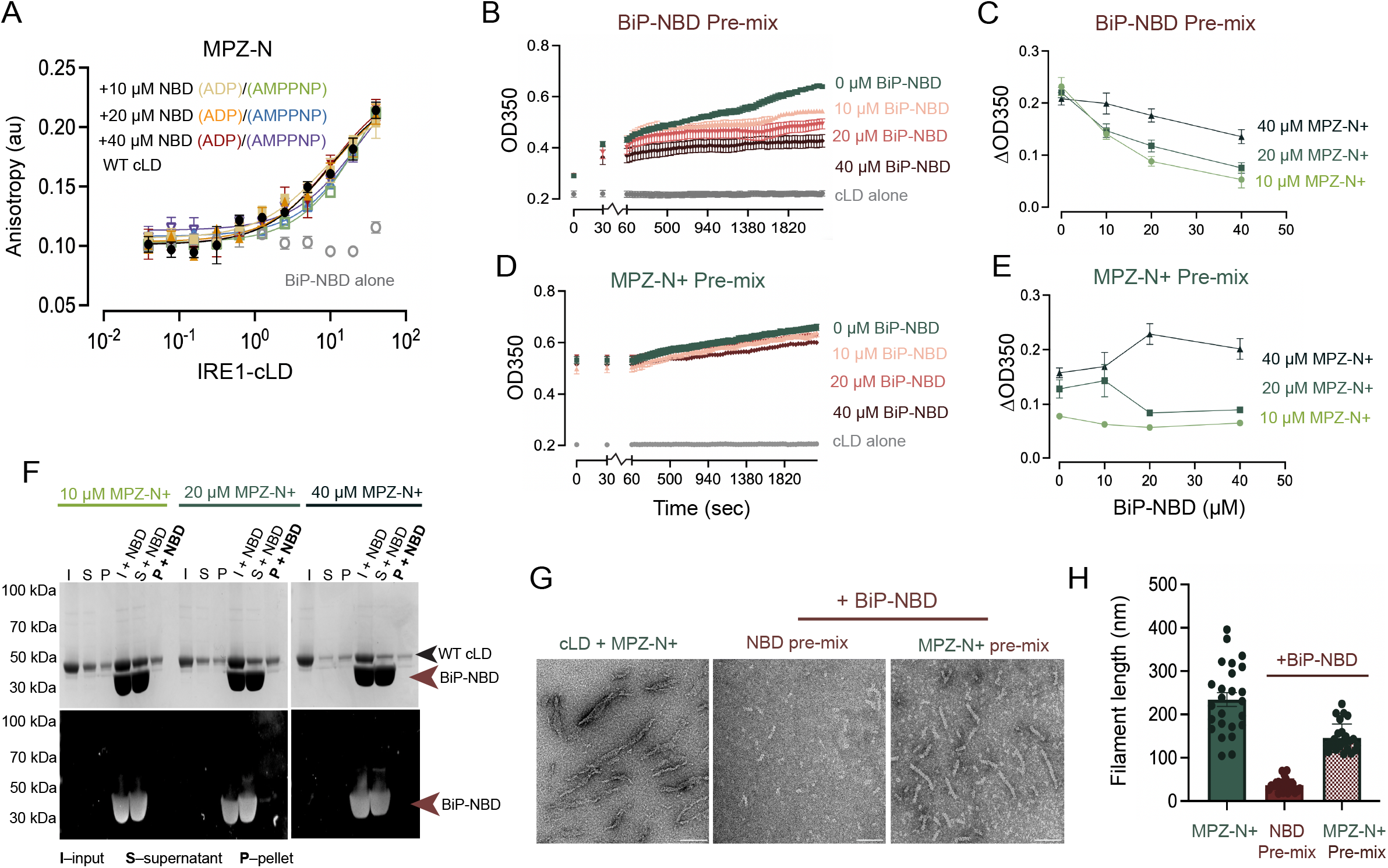
The BiP NBD impairs low-grade IRE1-cLD oligomerization. A. Increased concentrations of BiP-NBD bound to either ADP or AMP-PNP do not impair binding of IRE1-cLD to MPZ-N as determined by fluorescence anisotropy. B. Light scattering at UV 350 nm of 5 µM IRE1-cLD pre-mixed with increasing concentrations of BiP-NBD (0-40 µM) immediately following the addition of 20 µM MPZ-N+. C. Comparison of ΔOD350 values, defined as OD350_t=36m_ – OD350_t=30s_, obtained from different MPZ-N+ concentrations upon addition to IRE1-cLD and BiP-NBD (supporting data in SuppFigure 4A). D. Light scattering at UV 350 nm of 5 µM IRE1-cLD pre-incubated with 20 µM MPZ-N+ immediately following the addition of increasing concentrations of BiP-NBD (0-40 µM). E. Comparison of ΔOD350 values obtained when IRE1-cLD filaments were pre-assembled with different MPZ-N+ concentrations and subsequently challenged with increasing concentrations of BiP-NBD (supporting data found in SuppFigure 4B). F. SDS-PAGE and anti-BiP Western blots of pre-assembled IRE1cLD•MPZ-N+ filaments incubated with 40 µM BiP-NBD and separated by centrifugation into supernatant and pellet fractions. BiP-NBD is excluded from the pellet fraction containing IRE1 cLD filaments. G. Negative stain TEM micrographs (42kx mag and -1.04 µm defocus) of IRE1-cLD filaments induced by 20 µM MPZ-N+ compared to assemblies formed in the presence of BiP-NBD. H. Bar chart of IRE1-cLD filament lengths induced by MPZ-N+ in the presence or absence of BiP-NBD. Measurements of 25 filaments from each condition were performed using IMOD’s 3dmod program (79).

We next tracked light scattering of IRE1-cLD as a function of BiP-NBD concentration. To this end, we pre-mixed IRE1-cLD with varying amounts of BiP-NBD prior to the introduction of MPZ-N+ at a concentration shown to robustly induce filament formation (20 µM). Under these conditions, we observed that increasing concentrations of BiP-NBD (from 10-40 µM) dampened light scattering induced by MPZ-N+, indicating that the BiP-NBD impairs IRE1-cLD oligomerization (Fig. 4B). Accordingly, negative stain EM showed reduced filament assembly, indicated by shorter filaments, when MPZ-N+ was added to pre-mixed IRE1-cLD and BiP-NBD (Fig. 4G, H). Interestingly, the reduction in light scattering was partially suppressed by further increasing the MPZ-N+ concentration (to 40 µM) (Fig. 4C, SuppFig. 4B), suggesting that enhancing the speed of oligomerization expected to be induced by high MPZ-N+ may counteract the effect of BiP-NBD.

Consistent with this notion, when we pre-assembled filaments by incubating IRE1-cLD with MPZ-N+ prior to BiP-NBD addition, we observed no discernable effect on light scattered by the reaction (Figure 4D, E), and filaments were of comparable size with and without BiP-NBD. Filaments pelleted by centrifugation also did not incorporate BiP-NBD (Fig. 4F), indicating that filament assembly and BiP-NBD binding are mutually exclusive. Taken together, these results suggest that the BiP-NBD “buffers” the assembly of IRE1-cLD filaments, but fails to promote their disassembly.

Previously published reports showed that BiP-NBD interacts with IRE1-cLD monomers (35). As shown above, however, in our reaction, IRE1-cLD by itself was a dimer (SuppFig. 1A), which then oligomerized into filaments as observed. To reconcile these observations, we adapted a FRET pair sensitive to IRE1-cLD dimerization (29) by site-specific labelling of a single cysteine at IF1 with an Alexa Fluor 546-647 FRET pair. As expected, the FRET signal decreased when the labelled pair was incubated with increasing concentrations of unlabeled IRE1-cLD, indicating exchange between unlabeled and FRET IRE1-cLD monomers. Conversely, unlabeled cLD-ΔIF1 was unable to promote monomer exchange and did not result in FRET signal decrease (SuppFig. 4C), indicating that the assay reliably reports on the formation of IRE1-cLD dimers assembling via IF1.

When we titrated BiP-NBD into our FRET-labelled IRE1-cLD, we observed a concentration-dependent decrease in FRET signal, demonstrating that BiP-NBD binding to IRE1-cLD indeed destabilizes IF1 (SuppFig. 4D). We confirmed this conclusion by native MS, showing that the BiP-NBD forms a 1:1 stoichiometric complex with IRE1-cLD and affects an increase in the population of IRE1-cLD monomers relative to other species present (SuppFig. 4E,F).

### BiP-NBD binds to an unstructured loop at Interface 2 to drive allosteric inhibition of IRE1-cLD oligomerization

To map the binding interface of BiP-NBD and IRE1-cLD, we employed chemical crosslinking coupled to MS (XL-MS). We incubated IRE1-cLD and BiP-NBD with PhoX, a disuccinimidyl-based amine-selective crosslinker that allows selective enrichment of crosslinked products by metal-affinity chromatography. SDS-PAGE confirmed crosslinking products at the expected molecular mass (SuppFig. 5A). MS analysis identified several cLD to NBD lysine-to-lysine crosslinks that predominantly localized to Lobe IIA of the BiP NBD (Fig. 5A, thresholding score >40). Crosslink positions on IRE1-cLD were localized to two loops at IF2 (Fig. 5A). Both loops were unstructured in the crystal structure of human IRE1-LD dimers (25) and implicated in IRE1 oligomerization (26, 48).

**Figure 5.**
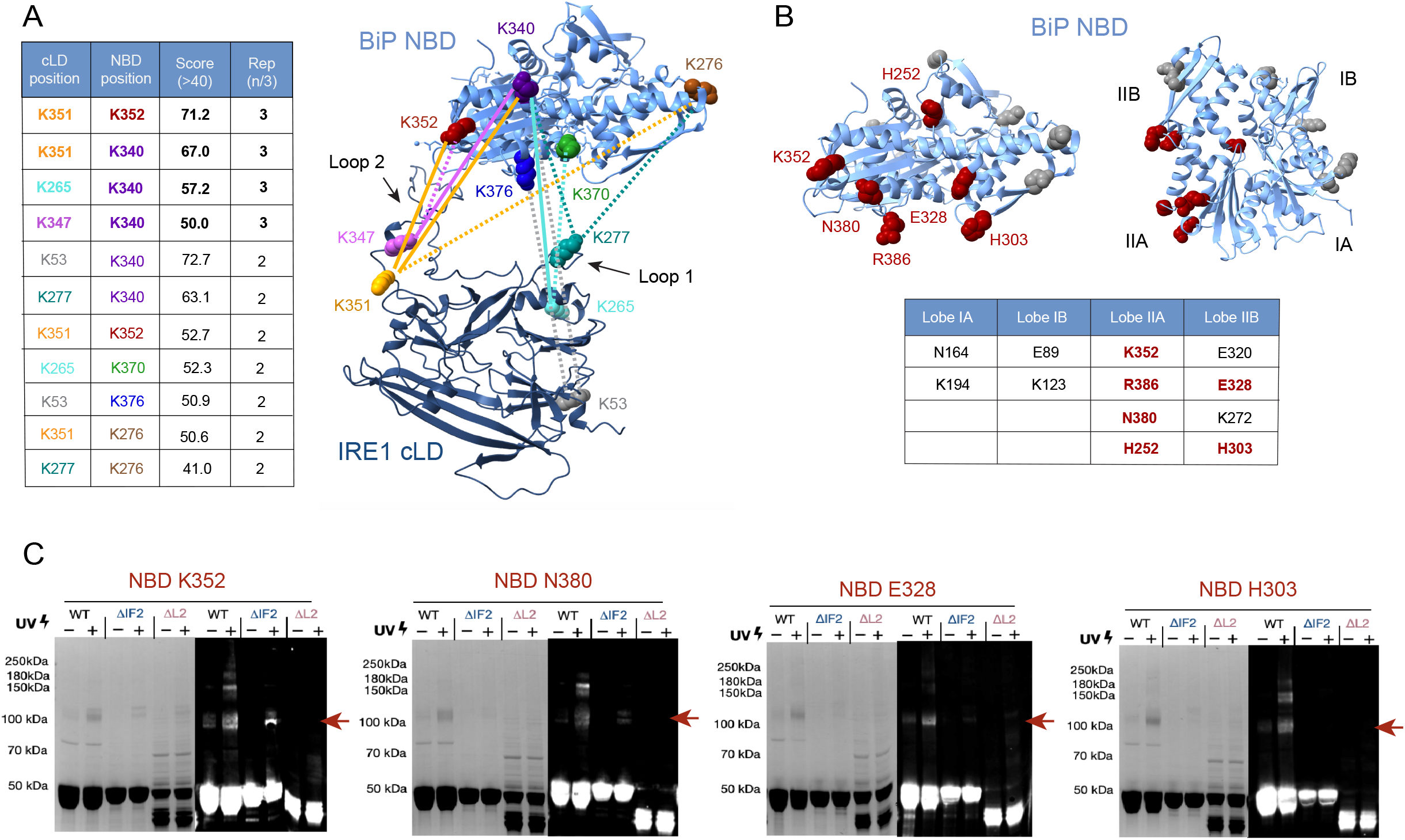
BiP NBD binding is localized to an unstructured loop at IRE1’s oligomerization interface. A. Table of cross-link sites identified by mass spectrometry mapped onto structural models of BiP-NBD and IRE1-cLD generated using AlphaFold3 (80) to provide better visualization of disordered regions. The mean confidence score and replication number (out of 3 independent sample replicates) for each cross-link site are reported. High confidence cross-links identified in all three sample replicates are indicated by bold lines and text (see Table 1). B. Positive IRE1-cLD photo-crosslink sites for p-Bpa insertion into 12 positions of BiP-NBD are mapped onto a BiP-NBD structural model generated using AlphaFold 3 (80). p-Bpa insertion sites by region of BiP-NBD are noted. Positively scored hits are localised to the Lobe IIA and Lobe IIB regions. C. SDS-PAGE gels and anti-IRE1 Western blots of photo-crosslinking observed between equimolar concentrations (10 µM) of known positive BiP-NBD hits (K352, N380, E328, H303) and IRE1-cLD variants. In all cases, truncation of the loop 2 region abolished cross-linking to BiP-NBD.

As orthogonal validation, we incorporated the near zero-length photoactivatable crosslinker p-benzoyl-L-phenylalanine (Bpa) at 12 positions distributed over the surface of BiP-NBD by incorporating amber codon (TAG) substitutions to insert the non-natural amino acid Bpa (49). We expressed the mutant proteins in *E. coli* and, after mixing with IRE1-cLD, exposed the mixture to UV light. We then scored the presence of IRE1-cLD containing crosslink products by Western blotting using anti-IRE1 antibodies (Fig. 5B). Six of the 12 mutants yielded robust crosslinks, indicated by arrows. Four of the six mapped to Lobe IIA, validating the MS analysis described above. The remaining two mutants that scored positively mapped to the adjacent Lobe IIB. We note that of the native lysines in Lobe IIB, six are located distally from the patch of amino acids identified on Lobe IIA and another two face towards the nucleotide binding groove, explaining why the XL-MS analyses did not precisely identify contributions of Lobe IIB to the IRE1-cLD binding site.

To further validate the role of the IF2 loops in binding to the BiP-NBD in this assay, we purified a truncation mutant in which the C-terminal loop was deleted (“ΔLoop2”; Δaa308-357). We picked four NBD-Bpa mutants (two in Lobe IIA, and two in Lobe IIB) that scored positively for cLD-binding, mixed them with either WT, ΔIF2, or ΔLoop2 cLD variants and performed UV-crosslinking. Notably, all four Bpa mutants failed to crosslink to ΔLoop2 cLD despite showing strong bands for the WT variant, underscoring that Loop 2 is essential for NBD binding (Fig. 5C). Two of the four (Bpa positions E328 and H303) also failed to crosslink with cLD-ΔIF2, suggesting that the IRE1-cLD•BiP-NBD interface is at least partially overlapping with IF2 involved in cLD oligomerization.

The binding site of BiP-NBD maps to the face of IRE1-cLD opposite to the amino acids involved in IF1 formation. Therefore, taken together, our data suggest a model in which the observed BiP-NBD mediated dissociation of IRE1-cLD dimers involves allosteric communication between IF1 and IF2, and the buffering effect of NBD on filament assembly involves sterically blocking IF2 and allosterically destabilizing the IRE1 dimer interface (IF1).

### IRE1-cLD filament disassembly requires a J domain co-chaperone and ATP hydrolysis

Since we showed that NBD-binding to IRE1-cLD monomers cannot resolve filaments once formed, we next asked whether filament disassembly is a mechanistically distinct process requiring energy input. To this end, we explored the canonical ATP-cycling capacity of BiP for filament disassembly. BiP is a poor ATPase on its own and requires a J domain co-chaperone to stimulate ATP hydrolysis as a prerequisite to reversible binding to clients (50-52). We challenged pre-assembled IRE1-cLD•MPZ-N+ filaments with full-length BiP and the ubiquitously expressed co-chaperone ERDJ3 (endoplasmic reticulum DnaJ protein 3) (53-55). Upon addition of ATP, we observed a decrease in light scattering (Fig. 6A) that correlated with shortened filament lengths in negative stain EM images (Fig. 6D,E), indicating that BiP and ERDJ3 actively dismantle IRE1-cLD filaments. The decrease in light scattering induced by BiP-ERDJ3 machinery was dependent on the presence of hydrolyzable ATP, as neither ADP nor AMP-PNP had an effect on the reaction (Fig. 6B). Moreover, it required an intact BiP SBD client binding pocket (Fig. 6B), confirming that filament disassembly occurs through BiP’s canonical ATP-regulated mechanism of client engagement.

**Figure 6.**
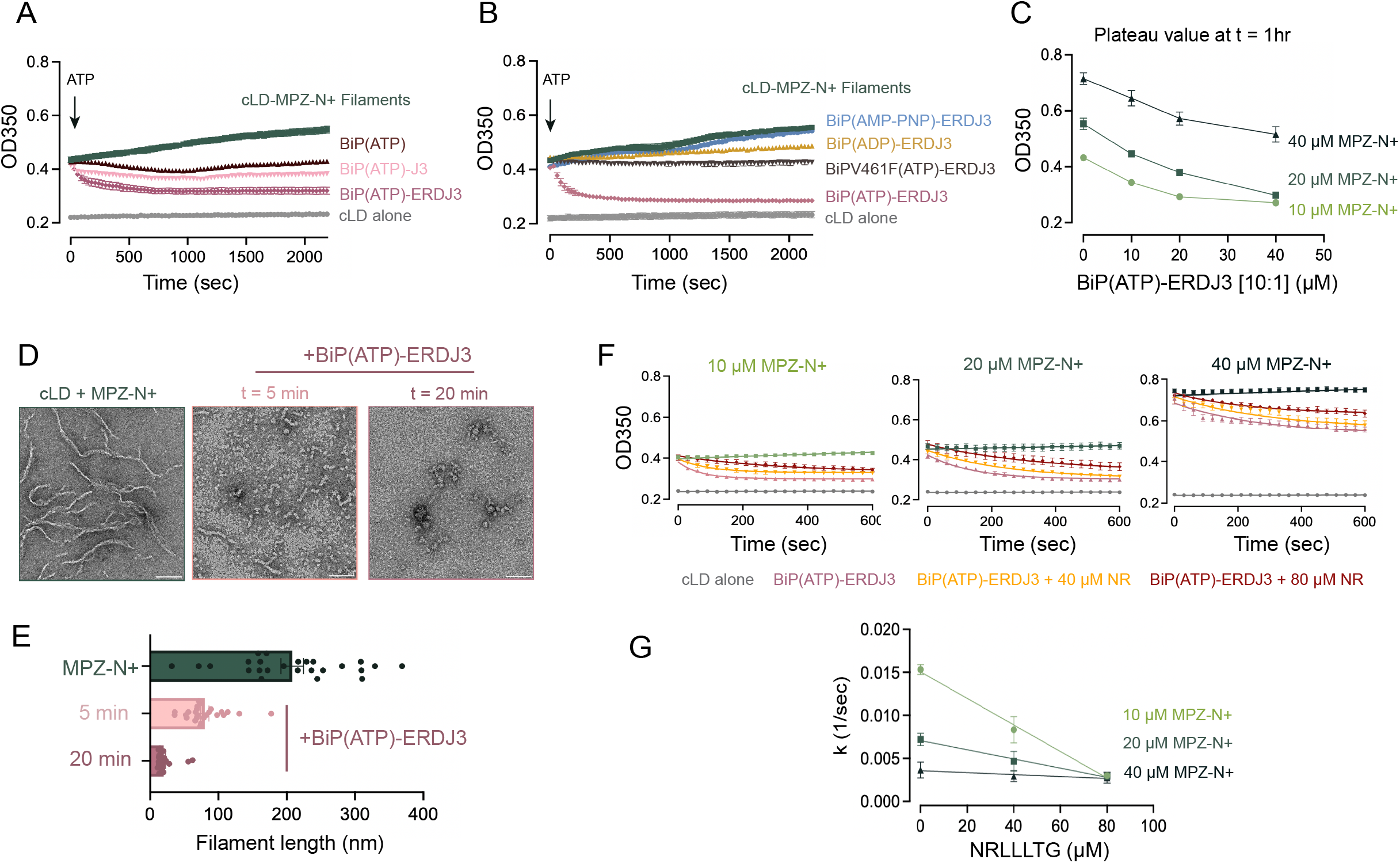
IRE1-cLD filament disassembly requires canonical chaperone-mediated ATP cycling. A. Light scattering at UV 350 nm of pre-assembled IRE1-cLD filaments (5 µM cLD + 20 µM MPZ-N+) in the presence of 10:1 full-length BiP(ATP) to ERDJ3. Only the complete chaperone machinery [BiP(ATP)-ERDJ3] decreases light scattering and mediates filament disassembly. B. IRE1-cLD filament (5 µM cLD + 20 µM MPZ-N+) disassembly by BiP/ERDJ3 requires hydrolyzable ATP and an intact substrate binding domain pocket. C. OD350 plateau (t = 1 hr) values for chaperone-mediated disassembly of IRE1-cLD filaments induced by different concentrations (10, 20, and 40 µM) of MPZ-N+ (support data found in SuppFig. 6). D. Negative stain TEM micrographs (42kx mag and -1.04 µm defocus) comparing IRE1-cLD filaments induced by MPZ-N+ to those present at different time points following the initiation of disassembly reactions with BiP(ATP)-ERDJ3. E. Bar chart of IRE1-cLD filament lengths induced by MPZ-N+ compared to those present at different time points following the initiation of disassembly reactions with BiP(ATP)-ERDJ3. Measurements of 25 filaments from each condition were performed using IMOD’s 3dmod program (79). F. IRE1-cLD filament (5 µM cLD + 20 µM MPZ-N+) disassembly by BiP(ATP)-ERDJ3 machinery in the presence and absence of NRLLLTG as measured by light scattering at 350 nm. G. Rate constants (k) of IRE1-cLD filament disassembly by BiP(ATP)-ERDJ3 in the presence of increasing concentrations (0, 40, 80 µM) of NRLLLTG as estimated from one phase exponential decay curve fittings performed in GraphPad Prism software.

By contrast, under the concentration and time regime used in the assay, ATP-bound BiP alone prevented further filament growth as assessed by a lack of further increase in light scattering but was insufficient to mediate disassembly (Fig. 6A). Including the isolated J domain of ERDJ3 (J3) did not lead to a decrease in light scattering (Fig. 6A), suggesting an impact on BiP’s ATP hydrolysis rate and/or dependence on ERDJ3’s known client-binding domains in promoting filament disassembly. A similar requirement of C-terminal ERDJ client-binding domains was observed in loading of the IRE1 lumenal domain onto BiP by a different ER DnaJ protein, ERDJ4 (29). Indeed, we found that ERDJ4 also disassembled IRE1-cLD filaments induced by MPZ-N+, and its isolated J domain was likewise insufficient (SuppFig. 6A). Therefore, our data show that multiple ERDJ proteins can stimulate filament disassembly.

We also explored the capacity of the BiP/ERDJ3 machinery to dismantle filaments induced by increasing concentrations of MPZ-N+. As previously shown, the extent of IRE1-cLD oligomerization, measured by light scattering, correlated with the amount of MPZ-N+ pre-incubated with IRE1-cLD to induce filament formation. In all cases, BiP and ERDJ3 mediated a concentration-dependent decrease in light scattering upon addition of ATP (Fig. 6C, SuppFig. 6B), indicating filament disassembly. However, when we examined the plateau of light scattering at 1 h for the highest concentration of BiP/ERDJ3 as a function of MPZ-N+ concentration, we found that filaments induced with higher concentrations of MPZ-N+ became progressively resistant to disassembly (Fig. 6C).

### The presence of an alternate BiP client impairs cLD filament disassembly by BiP/ERDJ3/ATP

Because IRE1-cLD filament disassembly requires BiP substrate engagement, we finally asked whether competition from an alternative BiP client would limit this process. To this end, we used NRLLLTG, a 7-amino-acid peptide that has been extensively used in mechanistic and structural studies of HSP70-type chaperones (56). Fluorescent 5FAM-NRLLLTG showed negligible binding to IRE1-cLD as measured by anisotropy (SuppFig. 6C), and NRLLLTG failed to induce IRE1-cLD filaments as measured by light scattering (SuppFig. 6D). When mixed in the absence of IRE1-cLD, the peptides did not scatter light, excluding the possibility that they engage in some unexpected aggregation that might confound the light scattering assay (SuppFig. 6E).

We next repeated the BiP/ERDJ3/ATP disassembly assay on pre-assembled IRE1-cLD•MPZ-N+ filaments in the presence of NRLLLTG added in a 1:1 or 1:2 molar ratio to BiP. As expected, NRLLLTG impaired the disassembly reaction at all MPZ-N+ concentrations used to preassemble the filaments (Fig. 6F, G). Disassembly rate constants (k) estimated by fitting light scattering curves to a one-phase decay model showed an inversely proportional relationship to NRLLLTG concentration (Fig. 6F), demonstrating that IRE1-cLD filaments can be shielded from disassembly when BiP becomes occupied with other clients.

## Discussion

The mechanism of unfolded protein recognition in the ER lumen leading to the activation of the UPR has been debated for well over two decades. For IRE1, two longstanding models have often been contrasted: i) direct activation by binding of unfolded proteins to its lumenal domain and ii) indirect activation by accumulating unfolded proteins titrating BiP inhibition away from IRE1. Here we dissect the role of key players in IRE1 activation and synthesize the complex network of interconnected processes that govern IRE1 activation into a unifying concept (Fig. 7). According to our model, in the absence of ER stress, IRE1 exists in multiple interconnected equilibria alternatively bound in reversible interactions to buffering factors, such as BiP (via its NBD), to itself in a dimer (linked through IF1), and to unfolded proteins (Fig. 7, Step 1, “Resting State”). Upon unfolded protein accumulation, this balance is shifted, increasingly populating the system with oligomeric IRE1 (linked through both IF1 and IF2) (Fig. 7, Step 3, “Active State”). Oligomerization of IRE1’s ER lumenal domain is the key step in IRE1 activation, bringing its cytosolic kinase and RNase domains into proper alignment to allow their trans-activation. Unfolded protein accumulation i) reduces the ER’s buffering capacity, e.g., of BiP by engaging it with its co-chaperones in its ATP-driven cycle and ii) directly binds to IRE1 LD thereby increasing its oligomerization propensity. Depending on the strength of ER stress, oligomers can but are not required to grow into filaments. Resolution of stress involves the catalytic disassembly of oligomers requiring full-length BiP, an Hsp40 and ATP-hydrolysis, actively resetting the system and restoring the steady state (Fig. 7, Step 4).

**Figure 7.**
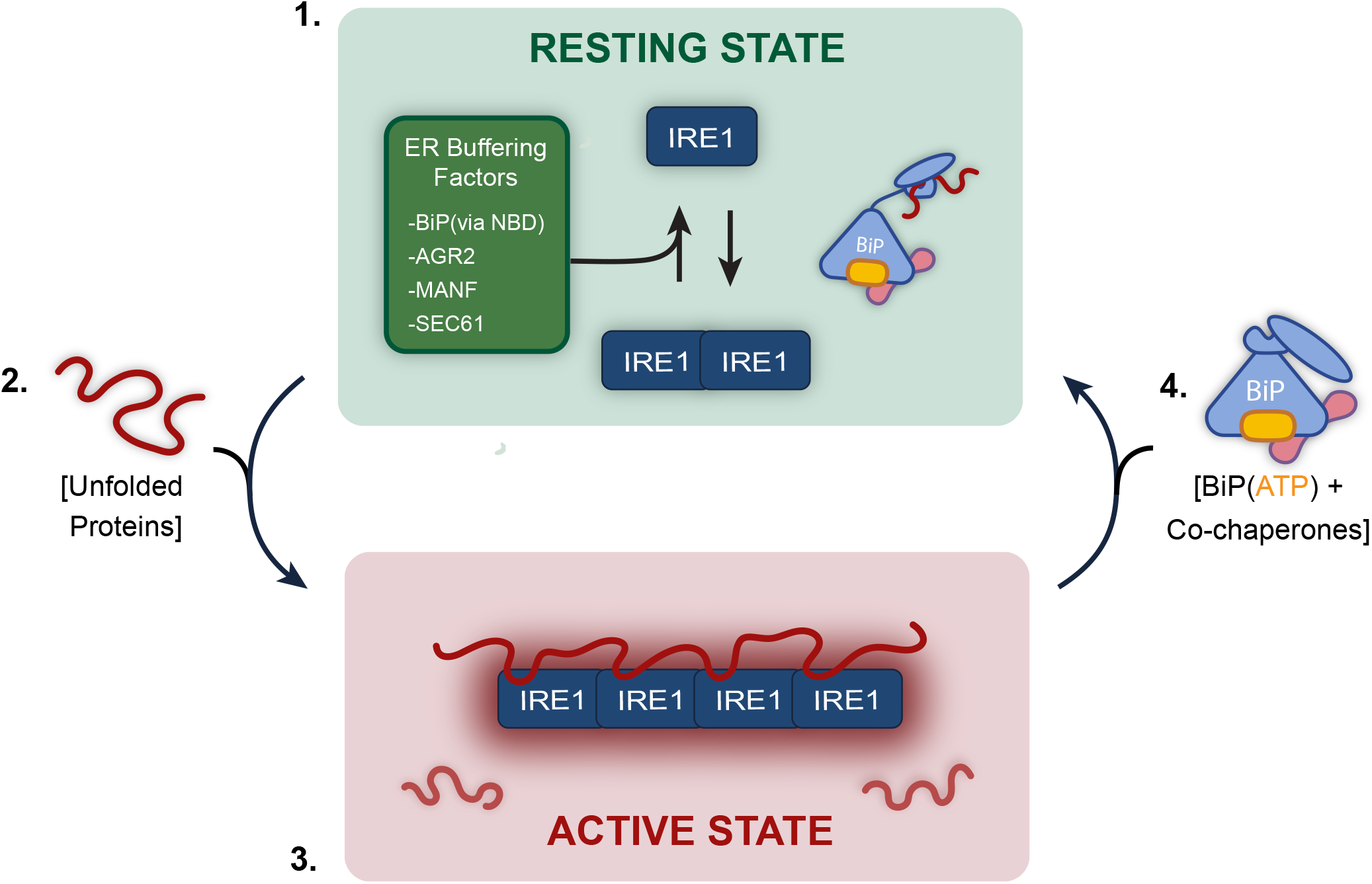
Model for dynamic regulation of IRE1 assembly by unfolded proteins and BiP. Step 1) In the absence of ER stress (“Resting State”), IRE1 is engaged in multiple interconnected processes that cumulatively bias it towards an equilibrium between two inactive (monomeric/dimeric) states through reversible and dynamic interactions with ER buffering factors/chaperones. **Step 2)** During the onset of stress, unfolded proteins accumulate in the ER, which reduce the buffering capacity of the ER lumen by engaging BiP in its ATP-driven cycle and bind IRE1. **Step 3)** In the “Active State”, the binding of unfolded proteins to IRE1 shifts the equilibrium towards an increasingly oligomeric state, which transmits stress detection from the ER lumen and activates IRE1’s cytosolic kinase and RNase modules. **Step 4)** Attenuation of IRE1 activity is triggered by a reduction in unfolded protein load, allowing for disassembly of IRE1 oligomers by energy-dependent BiP/ERDJ machinery.

Our work underscores that IRE1-cLD is a highly allosteric module, as previously proposed (26), in which IF1 and IF2, both required for IRE1 oligomerization, can be selectively switched on and off by interactions with other effectors binding to it. As such, unfolded proteins, BiP via its NBD, BiP via engagement of its ATPase cycle, and other buffering factors bind to different sites in IRE1-cLD and affect its oligomerization propensity.

### IRE1 buffering

According to our model, BiP is a dual-arm regulator of IRE1 oligomerization: passively via its NBD in addition to the known ATP-driven cycle of client engagement. Our observations support a role for the BiP-NBD in buffering IRE1 oligomerization. This is achieved through interaction with an unstructured loop at IRE1-cLD’s IF2, positioning the NBD to sterically interfere with oligomerization and allosterically destabilise dimerization via IF1. Indeed, overexpression of just the BiP-NBD reduces IRE1 activation in ER-stressed cells (35), indicating a less responsive UPR. The decrease in IRE1 activation results from the over-expressed pool of BiP-NBD, that is entirely de-coupled from its client engagement cycle, hence increasing buffering capacity. Although BiP is known to be a high abundance protein in the ER, a significant portion is sequestered in inactive oligomers (57) or is covalently inactivated (58, 59) in an unstressed cell, making it difficult to determine the true buffering capacity afforded by it at Resting State. However, the two main inactivating modifications, AMP-ylation and ADP-ribosylation, occur on known amino acids of BiP-SBD. Such modifications are unlikely to interfere with BiP-NBD binding to IRE1, strongly suggesting that even inactive “reserve” BiP contributes to Resting State buffering of IRE1 oligomerization.

In addition to BiP, cells contain other buffering factors that differ in abundance between cell types and tissues. AGR2, a protein disulfide isomerase expressed in mucin-producing goblet cells, destabilizes dimers of IRE1-beta (the IRE1-alpha paralog expressed in these cells) and represses its activity (60). As a monomer, AGR2 binds to loop 2 in IF2, thus competing with BiP NBD. Similarly, MANF, mesencephalic astrocyte-derived factor, represses IRE1 phosphorylation/splicing and reduces IRE1 oligomerization in cells by a yet unknown mechanism (61). An inhibitory association between IRE1 and the SEC61 translocon provides another layer of IRE1 surveillance sensitive to co-translational protein translocation (TRES) into the ER lumen (41, 62). Together, these factors maintain the resting equilibrium of IRE1 monomers and dimers (Step 1), while allowing the UPR to be tuned according to the physiological demands of different cell types and protein-folding environments.

Our biochemical data are consistent with the *in cellula* observation that a fraction of IRE1 molecules in unstressed cells assemble into inactive dimers via IF1 (20). BiP-NBD binding at IF2 and cLD homodimerization at IF1 are allosterically coupled but likely exist in a dynamic equilibrium of different inactive states. Through frequent transient associations with the cLD, excess BiP thus pushes IRE1 away from larger oligomers, tetramers and beyond. Thus, BiP-NBD binding at IF2 both destabilizes cLD dimers and acts as a competitive inhibitor of further cLD self-association past the dimeric state. This interpretation dovetails with previous cell culture studies showing that IF2 is strictly required both for stress-induced activation of IRE1’s cytosolic domains (26) and for the assembly of inactive IRE1 dimers into larger oligomers (20).

### IRE1 activation

As unfolded proteins begin to accumulate at the onset of ER stress, two events drive IRE1 towards a progressively more oligomeric state:

First, in the ER lumen the buffering imposed by BiP and other interactors is overcome, which frees up IRE1-cLD oligomerization interfaces. The engagement of BiP in its ATP-hydrolysis cycle by ERDJ3 is incompatible with IRE1-cLD binding to the BiP-NBD which impairs ERDJ3’s stimulation of BiP ATP-hydrolysis (37). Activation of BiP through removal of inactivating modifications further funnels existing chaperones towards populating the ATP-driven cycle (63). Similarly, the specialized collagen chaperone, HSP47, binds to IRE1-cLD and dislodges BiP (64), resulting in increased IRE1 oligomerization in ER-overloaded, collagen-producing cells. Thus, BiP engagement in ATP-dependent protein folding leads to a gradual loss in IRE1 buffering capacity in response to rising unfolded protein load (Step 2).

Second, IRE1 adopts an oligomerization-prone state through a direct interaction with unfolded proteins (Step 3). Our model proposes a direct effect of unfolded protein binding to IRE1-cLD on IRE1 activation via the assembly of lumenal domain oligomers and filaments. Model peptides reveal that unfolded proteins engage IRE1 monomers in a multivalent and IF2-dependent manner as a function of their length and sequence, as only long unfolded peptides with specific sequence elements induce assembly of IRE1-cLD into filaments. The IF2 dependence of binding by the “long” peptide (MPZ-N+) points to an extended interaction incorporating this interface, either directly or allosterically. This is consistent with observations of a distinct IRE1-cLD binding site employed by the A1 segment of cholera toxin (46) and is not in conflict with the notion that the “short” peptide (MPZ-N) interacts in the vicinity of the putative MHC-like groove as suggested by previous NMR and MD simulations (26, 65).

Yet the question of how unfolded proteins influence IRE1-cLD oligomerization propensity remains to be addressed. As previously reported, the short MPZ-N peptide elicits conformational rearrangements extending from the central beta-sheet floor (IF1) towards IF2 upon binding to IRE1-cLD (26). Indeed, IF2 has been identified as a hot spot of conformational heterogeneity through MD simulations (48), NMR spectroscopy (26) and, more recently, comparative AlphaFold studies (66). We consider it likely that binding of unfolded proteins is allosterically coupled to conformational rearrangements in IF2 that predispose IRE1-cLD to oligomerize (perhaps by exposing amino acids involved in IF2 or facilitating a restructuring of the interface), though an MPZ-N peptide bridging mode at IF2 cannot be fully excluded by our data. According to this view, these movements could be induced with a greater intensity and/or distance propagation along this trajectory by MPZ-N+, highlighting that larger UP domains engage in a more extended interaction with IRE1-cLD monomers that become increasingly coupled to IF2. Peptide-induced filament formation requires both IF1 and IF2. However, peptide binding still occurs at maximum affinity when IF1 is mutated, consistent with the notion that peptide binding within the monomer couples allosterically to IF2. We similarly observe allosteric coupling in IRE1’s interaction with the BiP-NBD, as NBD-binding to IF2 destabilized IRE1-cLD dimers at IF1, further underscoring the allosteric link between the two interfaces and effector binding sites.

In our assays, MPZ-N elicited only small oligomers in negative stain EM images, but proved insufficient for extended filament assembly elicited by longer peptides (MPZ-N+, PTIΔC). We attribute this enhancement in filament assembly to extended binding contacts that further stabilize the allosteric conformational rearrangements. This may explain, at least in part, why MPZ-N+ achieves extended filament formation at significantly lower peptide and IRE1-cLD concentrations compared to the 27-amino-acid ΔEspP peptide (66). Importantly, scrambled peptides of equivalent lengths did not induce filaments, indicating that functional binding contacts require specific features that remain to be defined. The question of what defines an IRE1-binding unfolded protein and what proportion of ER-processed proteins have this capability is critical for understanding the sensitivity of unfolded protein recognition by the UPR. Previous reports have suggested both structural motifs (46) and side chain preference for hydrophobics and positive charges (26) as a basis for ligand recognition.

Our observations directly tie unfolded protein binding to the assembly of IRE1 filaments similar to those previously observed in ER-stressed cells (67). Though IRE1-cLD oligomerization is generally accepted as the key step for IRE1 activation (17-20), the minimal oligomer unit sufficient for signaling remains unknown. Although large oligomeric structures may not be strictly required for signal activation (20, 68, 69), the degree of oligomerization can result in qualitatively different outputs, such as XBP1 mRNA splicing, RIDD, or RIDDLE, all involving catalytic engagement of IRE1’s RNase domain (70, 71). Additionally, they may allow IRE1 clusters to serve as a scaffold for downstream signaling effectors, such as TRAF2 (72). IRE1’s capacity for higher-order oligomerization is likely governed by its cellular concentration, which varies between cell types and disease states (11, 73), and the severity of a given proteotoxic insult. Though filaments likely represent extreme stress scenarios, the ability to monitor their assembly *in vitro* enables the unravelling of molecular principles underlying IRE1-cLD assembly.

### A role for disorder in IRE1 regulation

Growing evidence suggests that disordered regions of IRE1-cLD are important regulatory units for IRE1 signalling. Mutation/truncation of Loop 2 of the IRE1-cLD deregulates signaling in cells (30, 48). AGR2, mentioned above, binds Loop 2 of the epithelium-specific isoform, IRE1-beta, to destabilise IRE1 dimerization (60). Additionally, the A1 segment of cholera toxin was reported to bind a structural IRE1-cLD motif directly bordering Loop 2 (46). Hydrophobic patches in Loop 2 have even arisen as candidates for ATP-driven BiP engagement of IRE1-cLD (30, 66). Here, we expand on the importance of the disordered loop in showing that BiP NBD and unfolded protein ligands oppose each other’s action at this region. It is therefore interesting to speculate whether this unstructured loop might serve as a “regulatory hub” for IRE1 assembly where a range of interactors bias IRE1-cLD’s conformational heterogeneity or harness ATP-hydrolysis to direct a given outcome on IRE1 oligomerization propensity.

### IRE1 deactivation

UPR signaling drives adaptive mechanisms that reduce unfolded protein load in the ER and upregulate expression of ER chaperones. This achieves a decrease in unfolded protein load (ER stress), whilst adapting protein folding capacity. Together, these processes de-populate the system of oligomeric IRE1 by both removing the trigger for its oligomerization and actively dismantling existing oligomers (Fig. 7, Step 4). The disaggregation of disease-relevant amyloid fibrils (74, 75) and the disassembly of other transient oligomeric assemblies, such as clathrin coats, are well characterized for cytosolic Hsp70/Hsp40 machinery (76, 77). Our findings expand these observations into the lumen of the ER. IRE1 filament disassembly requires stimulated ATP-hydrolysis by an ERDJ co-chaperone, but is progressively stalled by stabilizing filaments with higher concentrations of unfolded proteins, indicating that elevated unfolded protein loads result in more persistent IRE1 oligomerization.

Resolving IRE1 oligomers therefore likely depends not only on direct disassembly, but also on general cellular strategies for reducing unfolded protein (i.e. chaperone-mediated refolding of proteins, ERAD) and protein folding load preemptively (RIDD/RIDDLE). Consistent with this notion, we observe that the presence of an alternate BiP client (NRLLLTG) slows IRE1 filament disassembly, suggesting that in unmitigated ER stress, BiP/ERDJ3 machinery may be occupied by other unfolded clients. A role for BiP/ERDJ machinery in oligomer disassembly is consistent with previous work underscoring the importance of BiP in IRE1 deactivation (32, 78) and with the *in cellula* observation that IRE1 molecules involved in signaling clusters can be recycled back into the ER network (69). These observations do not exclude the possibility of ERDJ-dependent monomerization of IRE1 at baseline (29); however, stress-induced upregulation of ER co-chaperones like ERDJ4 or handover by pre-existing SEC61/SEC63 (43) may serve to bias BiP engagement of IRE1 towards ATP-driven disassembly and thereby push the system to progressively lower levels of IRE1 oligomerization. The diversity of ERDJ family members and expression profiles may provide an additional layer of spatio-temporal control of IRE1 de-oligomerization.

## Conclusions

The model proposed here integrates many previously discussed concepts into a cohesive framework. Oligomerization of IRE1’s lumenal domain remains the key process that cells must continuously monitor and control for proper sensing of and response to ER stress. This complex task is orchestrated by a large number of ER-resident factors, some of which are general and present in every cell, whereas others are specialized to the unique requirements of particular cell types and their respective protein folding needs. BiP now emerges as an important regulator of two distinct processes: buffering and resetting, thus allowing the UPR to be tuned according to the physiological demands of the ER’s protein-folding environment. Rather than representing mutually exclusive mechanisms, direct unfolded protein recognition and BiP association can be understood as interconnected inputs whose balance promotes, buffers, and resolves IRE1 oligomerization, allowing for continuous adjustment of IRE1 signalling capacity emerging from dynamic control of its lumenal domain’s assembly state.

## Supporting information

Table1_IRE1BiP_PhoXCrosslinks

## Acknowledgements

We thank the current and former members of the Walter and Agard labs for their insightful comments and support throughout this work, particularly Advait Subramanian, Vlad Belyy, Chari Noddings, and Michael Schoof. We thank Johannes Buchner, Jason Gestwicki, and Kliment Verba for many helpful scientific discussions. Electron microscopy was performed with the support of UCSF EM facility managers David Bulkley, Glenn Gilbert, Li Wang, and Matt Harrington. The EM Facility at UCSF is supported by NIH grants (S10OD020054, S10OD021741 and S10OD026881) and the Howard and the Howard Hughes Medical Institute. Native MS was performed with the support of UCSF MS facility manager Mark Burlingame.

## Author Contributions

M.M-M., P.W., and D.A.A conceptualized the work, interpreted results, and wrote the manuscript. M.M-M. performed experiments, analyzed data, and wrote the original draft of the manuscript under the supervision of P.W. and D.A.A. M.L-V. and D.N.I performed cross-linking mass spectrometry sample prep, experiments, and analysis. W.S. trained M. M-M. in Native MS methodology and assisted in experiments/spectra deconvolution. G.E.K performed peptide microarray experiments during her time as a Walter lab member and shared analysed data with M. M-M. All authors read, edited, and approved the final version of the manuscript.

---

**SuppFigure 1.**
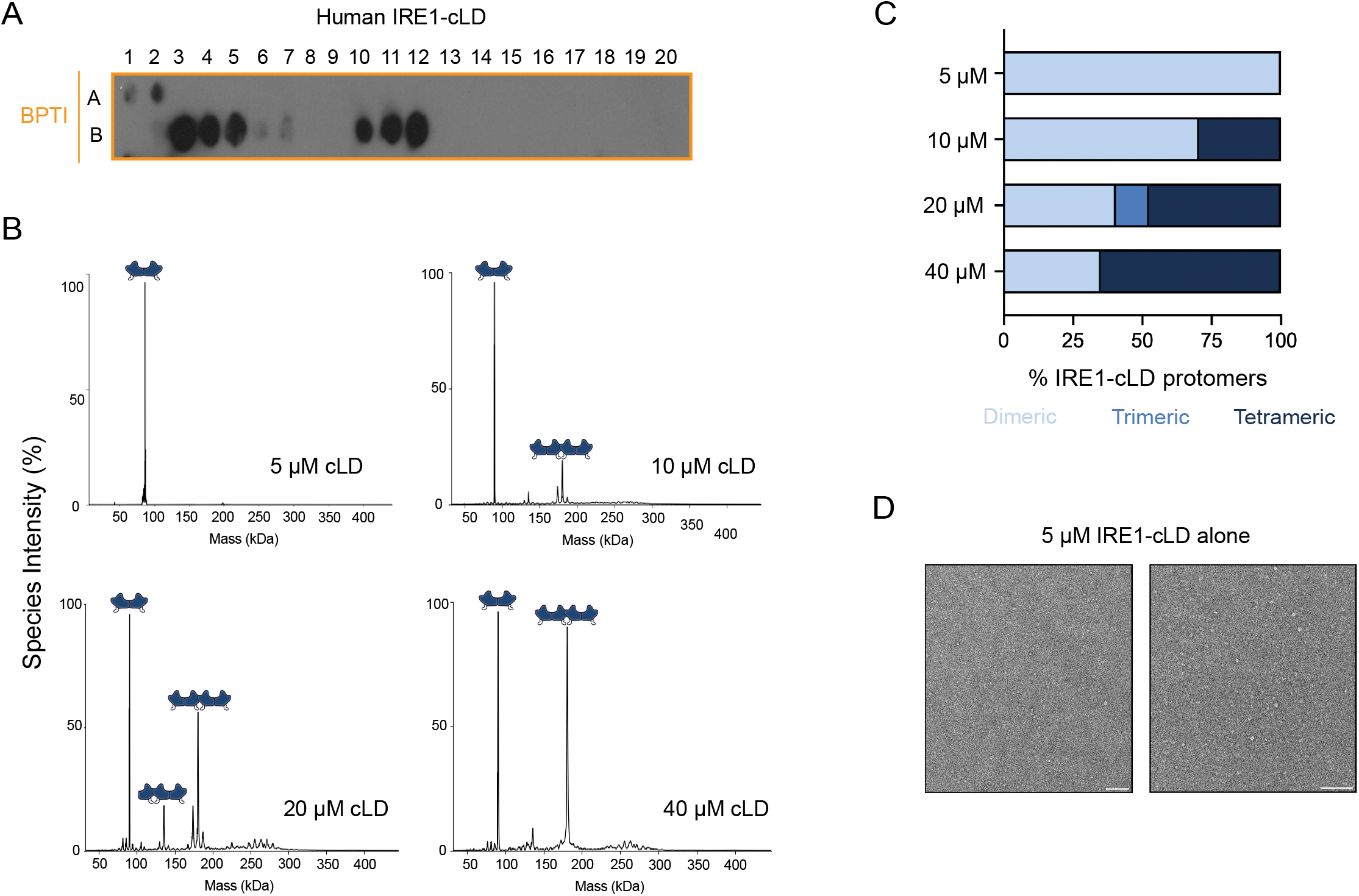
A. A peptide array tiled with 18mer peptides derived from bovine pancreatic trypsin inhibitors and probed with MBP-tagged human IRE1-cLD shows two regions of binding. B. Masses in kDa of the oligomeric states of IRE1-cLD at 5, 10, 20, and 40 µM present in solution as determined by native mass spectrometry. C. Bar chart displaying the percent of IRE1-cLD protomers making up each of the observed oligomeric states calculated from the experimental species intensities. IRE1-cLD oligomerization state in solution increases in a concentration-dependent manner. D. Negative stain TEM micrographs (42kx mag and -1.04 µm defocus) of 5 µM IRE1-cLD alone.

**SuppFigure 2.**
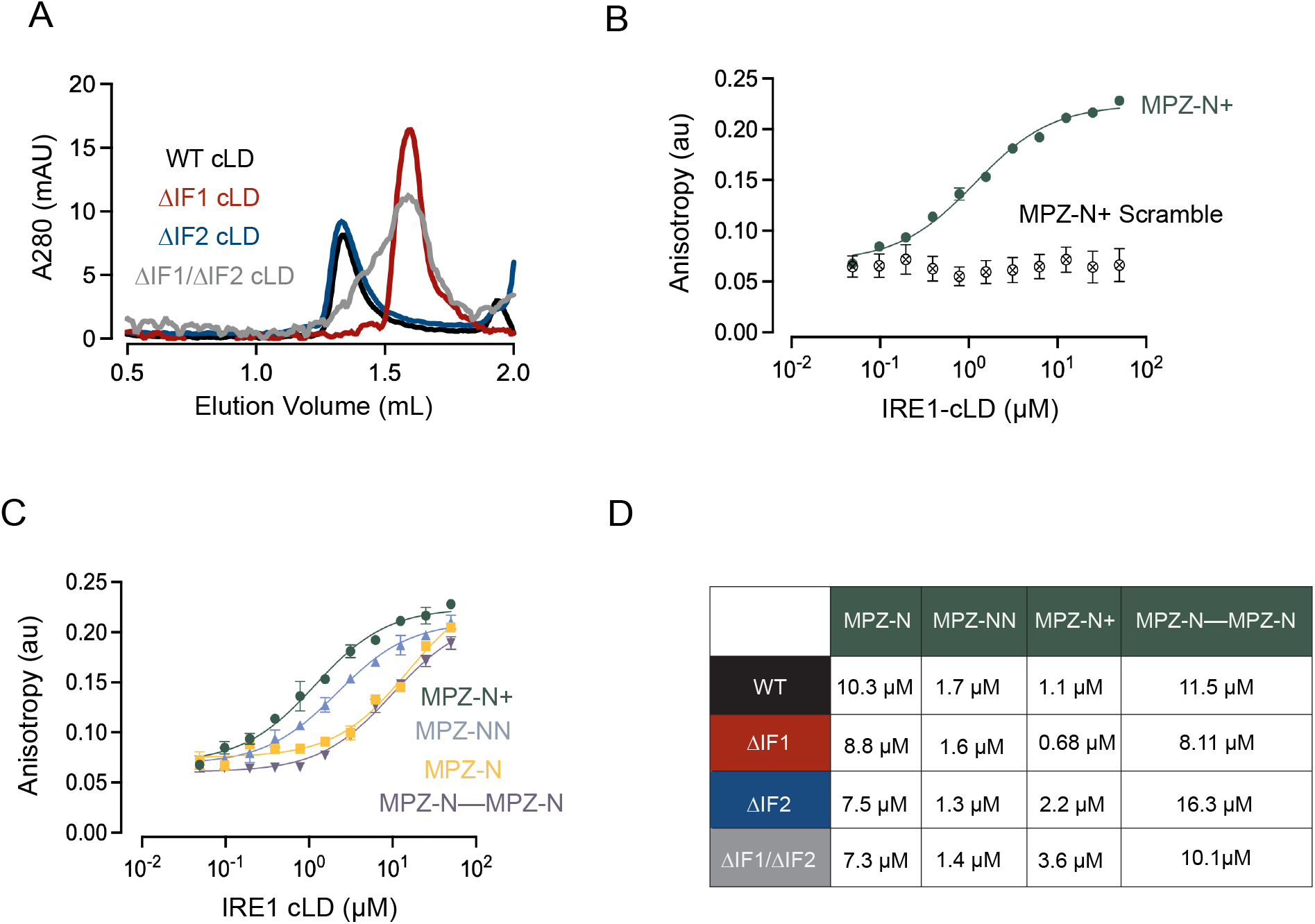
A. Size exclusion chromatography traces of 5 µM IRE1-cLD and its interface mutants run on an S200 3.2/300 column show that variants incapable of forming IF1 elute in later fractions. B. A scrambled sequence of MPZ-N+ (generated with an online scrambler tool at <u>peptidenexus.com</u>, sequence: YRAEALLGRPKSKPIMYVIDGRRRSLAMGLTSKSGQGRAWLAQKRLQH) shows no binding to IRE1-cLD in a fluorescence anisotropy assay. C. Comparison of IRE1-cLD binding to all MPZ-derived peptides used in Figure 2 as determined by fluorescence anisotropy. MPZ-N+ shows the highest affinity binding. D. Table of K_1/2_ values calculated from fluorescence anisotropy experiments binding IRE1-cLD and its interface mutants to MPZ peptides. Curve fittings were performed in GraphPad Prism software.

**SuppFigure 3.**
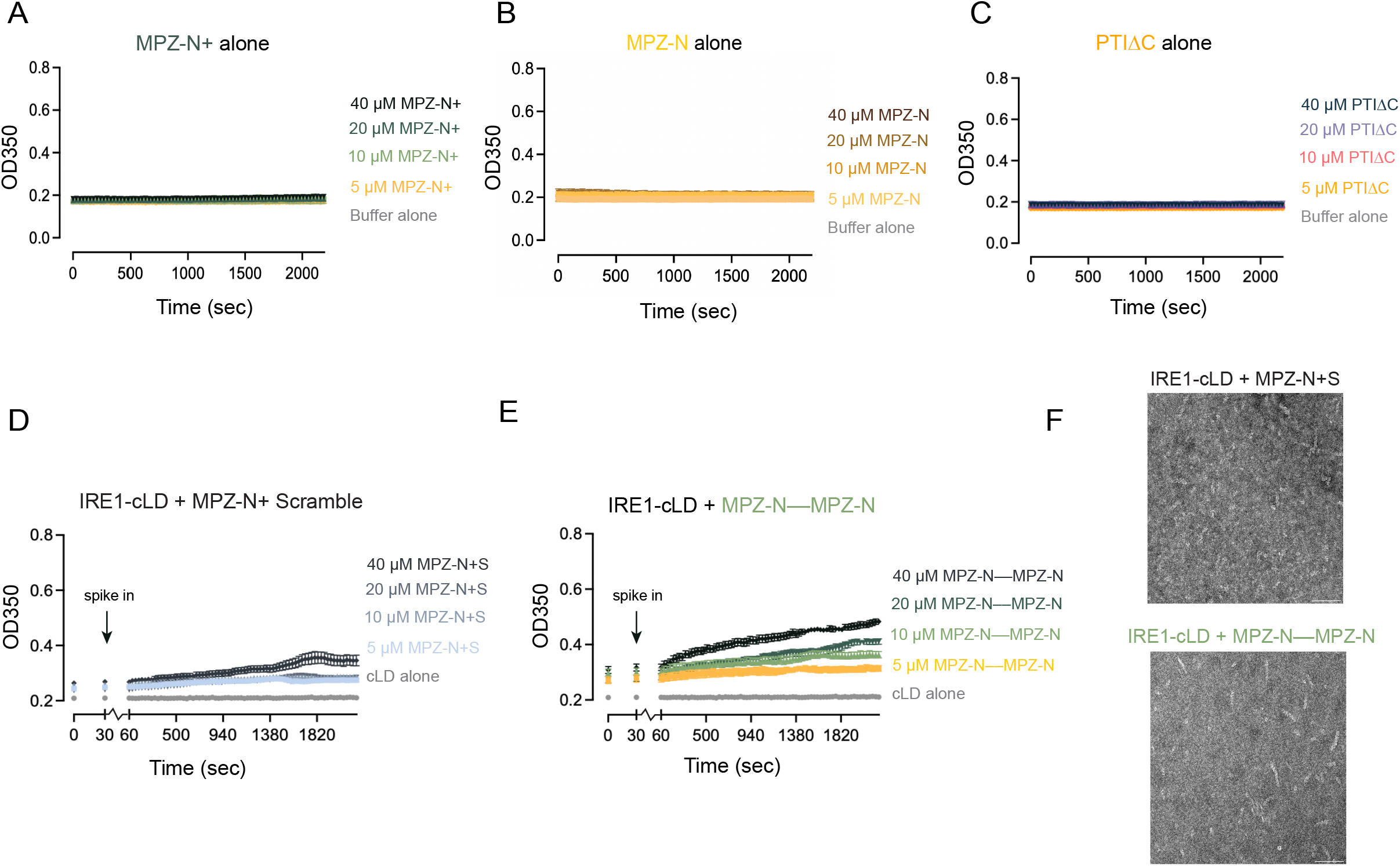
A. Light scattering at UV 350 nm of increasing concentrations (0-40 µM) of MPZ-N+ in solution over a 1 hr period at 25°C. B. Light scattering at UV 350 nm of increasing concentrations (0-40 µM) of MPZ-N in solution over a 1 hr period at 25°C. C. Light scattering at UV 350 nm of increasing concentrations (0-40 µM) of PTIΔC in solution over a 1 hr period at 25°C. D. Light scattering of 5 µM IRE1-cLD in the presence of increasing concentrations (0-40 µM) of the scrambled MPZ-N+ sequence (MPZ-N+S) over a 1 hr period at 25°C. E. Light scattering of 5 µM IRE1-cLD in the presence of increasing concentrations of the extended tandem binding site peptide MPZ-N––MPZ-N (0-40 µM) over a 1 hr period at 25°C. F. Negative stain TEM micrographs (42kx mag and -1.04 µm defocus) of WT IRE1-cLD or its variants in the presence of either the scrambled MPZ-N+ peptide or MPZ-N––MPZ-N.

**SuppFigure 4.**
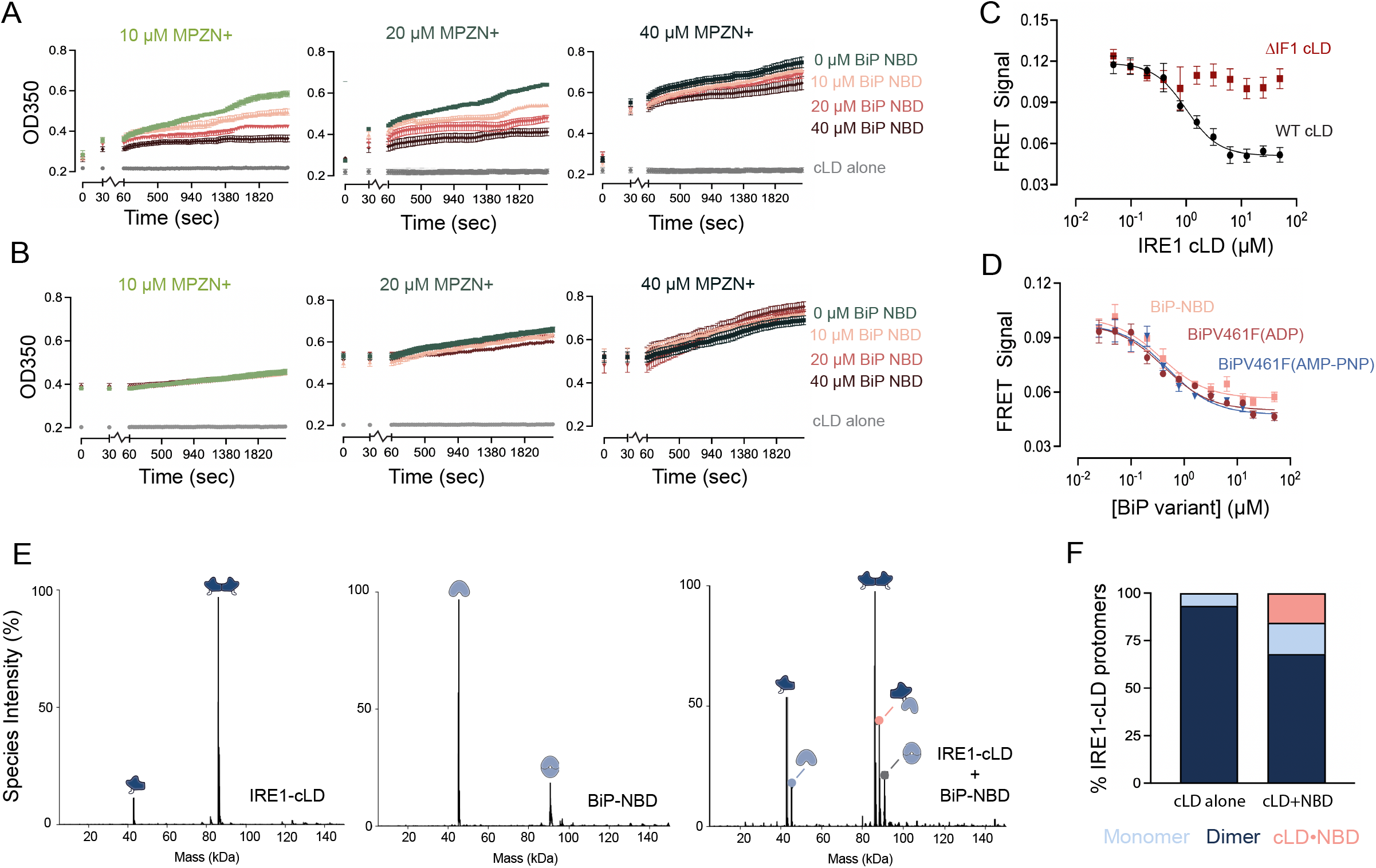
A. Light scattering at UV 350 nm of 5 µM IRE1-cLD pre-mixed with increasing concentrations of BiP-NBD (0-40 µM) immediately following the addition of the indicated concentration of MPZ-N+. B. Light scattering at UV 350 nm of 5 µM IRE1-cLD pre-incubated with 10, 20, or 40 µM MPZ-N+ immediately following the addition of increasing concentrations of BiP-NBD (0-40 µM). C. A cys-lite IRE1 cLD was site-specifically labelled by introduction of a single cysteine at position 112 for maleimide conjugation to either AlexaFluor-546 or AlexaFluor-647 dye. When titrated, unlabelled WT IRE1-cLD exchanges with FRET-labelled IRE1-cLD to decrease FRET signal (Em at 670nm/Em at 580nm) across IF1. D. Both BiP-NBD and full-length BiPV461F (ADP or AMP-PNP) mediate a decrease in FRET signal (Em670/580 values) across IF1. E. Masses in kDa of the species present in solution when 5 µM IRE1-cLD was incubated with 13 µM BiP-NBD as determined by native mass spectrometry. F. Bar chart displaying the percent of IRE1-cLD protomers present in each of the IRE1-containing species calculated from the experimental species intensities. The BiP-NBD forms a 1:1 hetero-complex with IRE1-cLD and decreases the amount of dimeric IRE1-cLD.

**SuppFigure 5.**
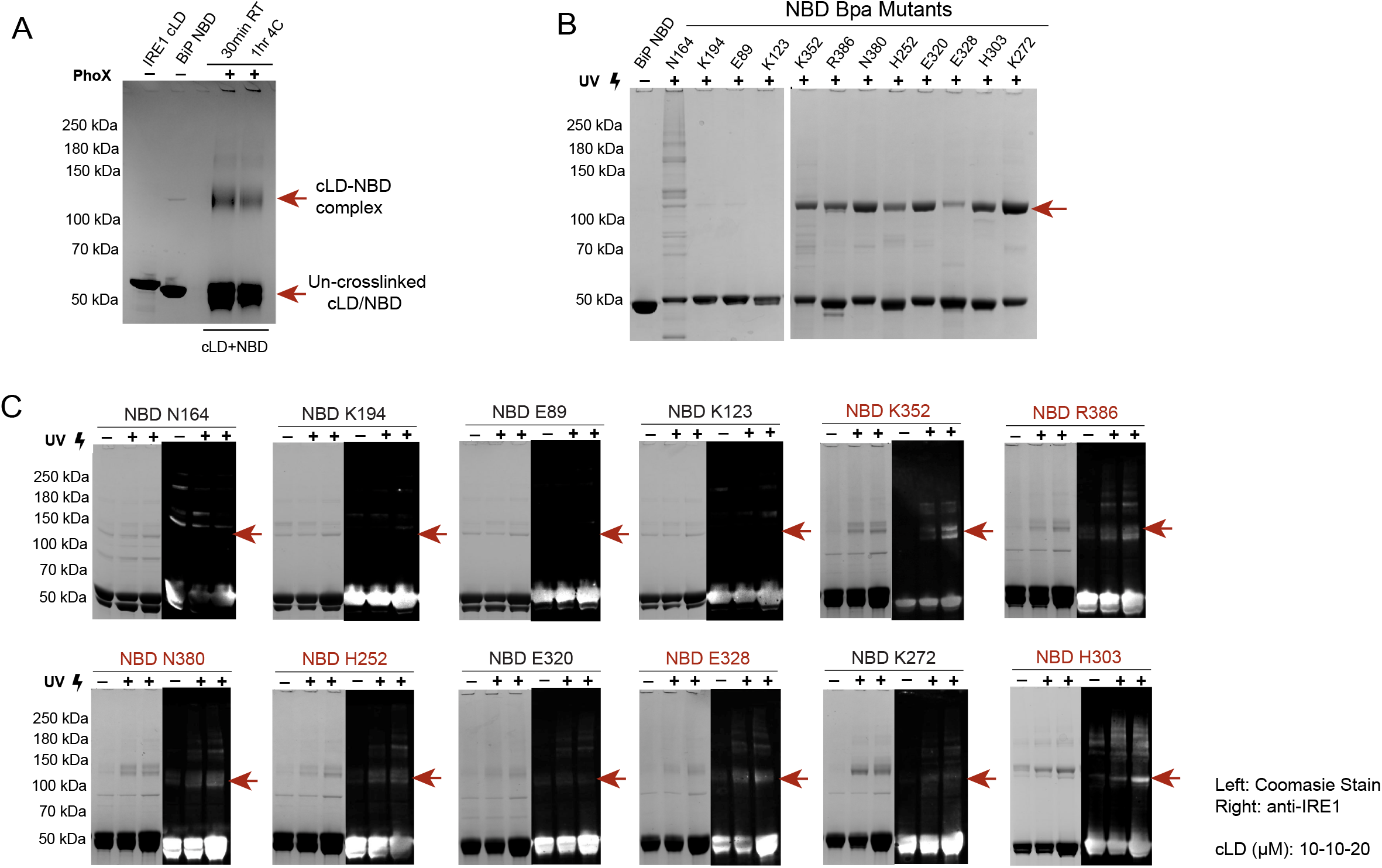
A. SDS-PAGE gel of equimolar (10 µM) IRE1 cLD and BiP NBD cross-linked by PhoX showing formation of an IRE1-cLD•BiP-NBD crosslink product species at higher MW. B. SDS-PAGE gel of 10 µM BiP-NBD or p-Bpa insertion mutants with or without exposure to UV light (265 nm) for 1 hr at 4°C. Some p-Bpa mutants show considerable self-crosslinking when alone. C. SDS-PAGE and anti-IRE1 Western blots of IRE1 cLD and all p-Bpa-insertion mutants (either 10:10 or 10:20 µM ratio) exposed to UV light (265 nm) for 1 hr at 4°C. Crosslink bands are indicated.

**SuppFigure 6.**
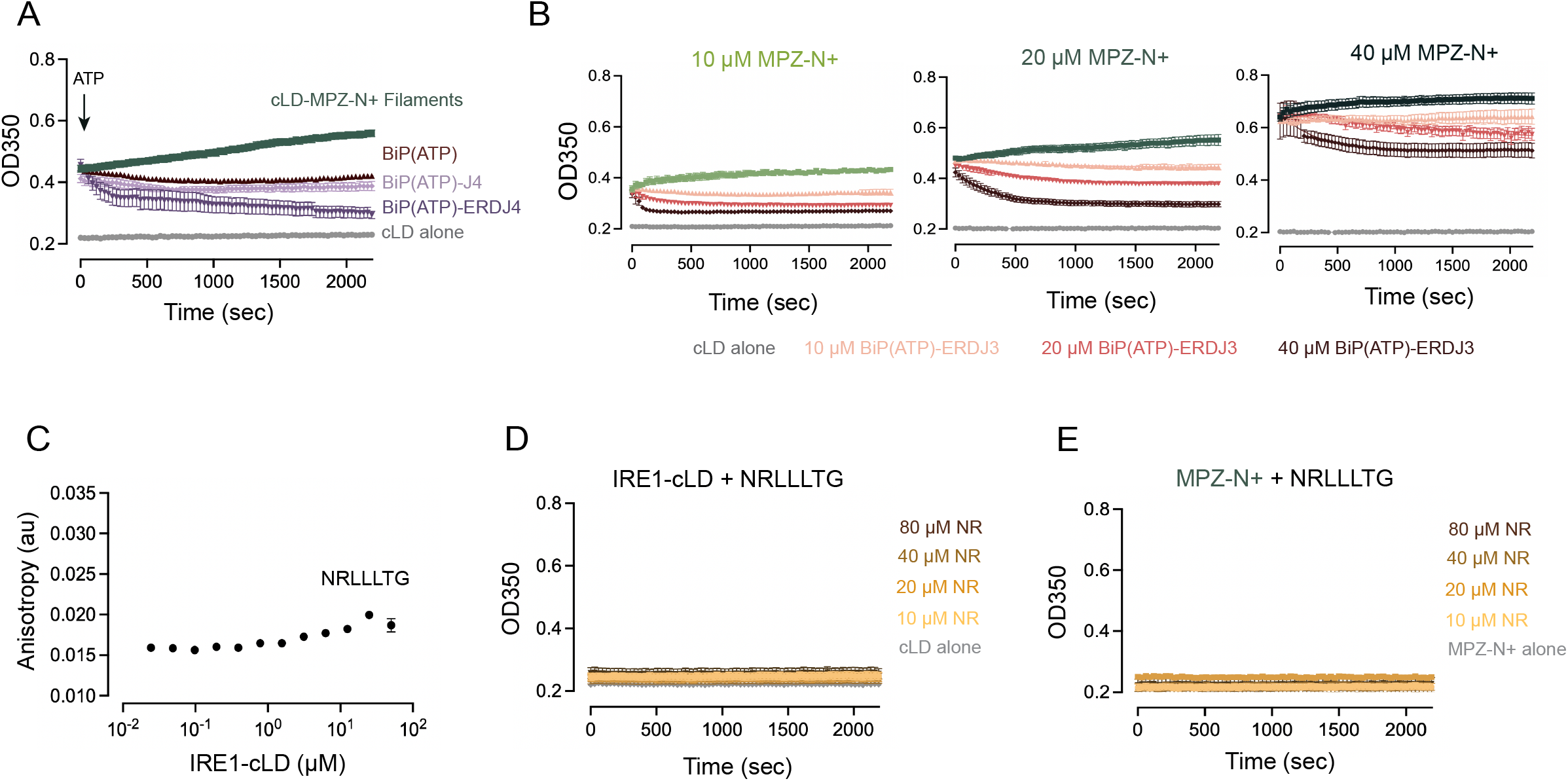
A. Light scattering at UV 350 nm shows that pre-assembled IRE1-cLD filaments (5 µM cLD + 20 µM MPZ-N+) can also be dismantled by BiP(ATP) and the stress-inducible co-chaperone ERDJ4. B. IRE1-cLD filament disassembly by increasing concentrations (0-40 µM) of BiP(ATP) and ERDJ3 at a 10:1 ratio. Filaments were pre-assembled with increasing concentrations (10, 20, and 40 µM) of MPZ-N+. C. IRE1-cLD shows no specific binding to NRLLLTG as measured by fluorescence anisotropy. D. Light scattering at UV 350 nm of 5 µM IRE1-cLD in the presence of increasing concentrations of NRLLLTG peptide (0-80 µM) over a 1 hr period at 25°C. E. Light scattering at UV 350 nm of 20 µM MPZ-N+ in the presence of increasing concentrations of NRLLLTG (0-80 µM) over a 1 hr period at 25°C.

## Materials and Methods

### Protein Expression and Purification

The plasmids and vectors used in this study are summarized in **Table 2**. For expression, the plasmid of interest was transformed into the BL21 DE3 strain of Escherichia coli (Agilent Technologies) unless otherwise specified. Cells were grown in LB with the appropriate antibiotic at 37°C until OD600 = 0.6. Expression was then induced with 0.4 mM IPTG for 18 hrs at 18°C.

**Table 2.** Summary of constructs and their vectors.

| Name | Description | Backbone | Construct Tags | Marker |
| --- | --- | --- | --- | --- |
| Hs_IRE1cLD | Human ERN1 (22-402) (cLD) | pet16b+ | N term 10xHis | AMP |
| Hs_IRE1cLD_ΔIF1 | Human ERN1 cLD K121Y | pet28a+ | N term 6xHis | KAN |
| Hs_IRE1cLD_ΔIF2 | Human ERN1 cLD 359-362 to GSGS | pet47b+ | N term 6xHis | KAN |
| Hs_IRE1cLD_ΔIF1/ΔIF2 | Human ERN1 cLD both ΔIF1 and ΔIF2 | pet28a+ | N term 6xHis | KAN |
| Hs_IRE1cLD_IF1FRET | Human ERN1 cLD cyslite and S112C | pet28a+ | N term 6xHis | KAN |
| Hs_IRE1cLD_Δloop2 | Human ERN1 cLD ΔP308-N357 | pet28a+ | N term 6xHis | KAN |
| MBP_Hs_IRE1cLD | Human ERN1<br>(22-402) (cLD) | pmalC2 | N term MBP | AMP |
| Hs_BiPNBD | Human HSPA5<br>(20-407) | pet29b+ | N term 6xHis | KAN |
| Hs_BiPFL | Human HSPA5<br>(28-654) | pet29a+ | N term 6xHis | KAN |
| Hs_BiPV461F | Human HSPA5<br>(28-654) V461F | pet29b+ | N term 6xHis | KAN |
| Hs_ERDJ3 | Human DNAJB11<br>(23-358) | pet28a+ | N term<br>6xHisSUMO | KAN |
| Hs_ERDJ3_Jdomain | Human DNAJB11<br>(23-96) | pet28a+ | N term<br>6xHisSUMO | KAN |
| Hs_ERDJ4 | Human DNAJB9<br>(24-223) | pet21a+ | N term<br>6xHisSUMO | AMP |

For human IRE1 cLD and human BiP variants, cells were resuspended in Lysis Buffer (50 mM HEPES pH 7.2, 400 mM KCl, 5 mM beta-mercaptoethanol (BME), 5% glycerol, and 2 mM Imidazole) and lysed by an Avestin Emulsiflex-C3 at 16,000 psi. Lysate was clarified by centrifugation at 30,000xg for 35 min. His-tag fusion constructs were nutated with equilibrated Ni-NTA resin in a gravity column for 1 hr at 4°C before washing 2X with Lysis Buffer and 2X with ATP Wash Buffer (50 mM HEPES pH 7.2, 200 mM KCl, 5 mM BME, 5 mM MgCl_2_, 5 mM ATP, and 2 mM Imidazole). Proteins were eluted in Elution Buffer (25 mM HEPES, 100 mM KCl, 300 mM Imidazole, and 5 mM BME). The eluate was further purified by anion exchange chromatography (MonoQ column) with MonoQ Buffers A (25 mM HEPES pH 7.2, 50 mM KCl, 5 mM BME, and 5% glycerol) and B (25 mM HEPES pH 7.2, 1 M KCl, 5 mM BME, and 5% glycerol) before His-tags were removed by either PreScission protease (1 unit enzyme per 100 µg protein) or FactorXa (1 µg enzyme per 100 µg protein) at 4°C overnight with protein concentration adjusted to 2 mg/mL. BiP constructs were dialysed against Dialysis Buffer (25 mM HEPES pH 7.2, 75 mM KCl, and 2 mM BME) concurrently with overnight tag cleavage to ensure removal of bound nucleotides. Separation of uncleaved protein and protease was performed by negative purification in Ni-NTA resin and the flow-through was loaded onto a size exclusion column (Superdex 200 16/600) equilibrated with SEC Buffer (25 mM HEPES pH 7.2 and 75 mM KCl). The eluate was pooled and concentrated in Cytiva Vivaspin columns before flash freezing in liquid nitrogen and storing at -80°C.

For human ERDJ3 and human ERDJ4 variants, constructs were transformed into Origami B DE3 cells (Novagen) to better preserve native disulphide bond formation and grown as previously described. Cells were resuspended in non-reducing (N-R) Lysis Buffer (50 mM HEPES pH 7.2, 400 mM KCl, 5% glycerol, and 2 mM Imidazole) and lysed by Emulsiflex. Lysate was clarified via centrifugation at 30,000xg for 35 min and the supernatant was incubated with pre-equilibrated Ni-NTA resin in a gravity column for 1 hr at 4°C. Four washes were performed: washing 2X with N-R Lysis Buffer and 2X with N-R ATP Wash Buffer (50 mM HEPES pH 7.2, 200 mM KCl, 5 mM MgCl_2_, 5 mM ATP, and 2 mM Imidazole). Proteins were eluted in N-R Elution Buffer (25 mM HEPES, 100 mM KCl, and 300 mM Imidazole) and loaded onto a size exclusion column (Superdex 200 10/300) equilibrated with SEC Buffer. Fractions containing ERDJ proteins were collected, flash frozen, and stored as described above.

### IRE1 and BiP Bpa Mutant Expression

For expression of benzoyl-phenylalanine (Bpa) incorporated mutants of human IRE1 cLD and human BiP (**Table 3**), plasmids were transformed into BL21 DE3 cells already containing a pEVOL-pBpF plasmid as described by Chin et al. (49). Cells were grown in TB with the appropriate double antibiotic markers at 37°C until OD = 0.2. Expression and loading of the amber suppressor tRNA were induced by the addition of 0.02% arabinose and 1 mM Bpa. Cells were allowed to continue growing at 37°C until OD = 0.8, when expression of the protein of interest was induced by 0.4 mM IPTG for 18 hrs at 16°C. Purification was performed as described above for BiP variants with samples being kept in the dark as much as possible.

**Table 3.** Summary of Bpa-incorporated BiP NBD mutants.

| Name | BiP Region | Backbone | Expression Line | Markers |
| --- | --- | --- | --- | --- |
| Hs_BiPNBD_N164* | Lobe 1A | pet28a+ | BL21 DE3 +<br>pEVOL-pBpF plasmid | KAN/CAM |
| Hs_BiPNBD_K194* | Lobe 1A | pet28a+ | BL21 DE3 +<br>pEVOL-pBpF plasmid | KAN/CAM |
| Hs_BiPNBD_D418* | Lobe 1A | pet28a+ | BL21 DE3 +<br>pEVOL-pBpF plasmid | KAN/CAM |
| Hs_BiPNBD_E89* | Lobe 1B | pet28a+ | BL21 DE3 +<br>pEVOL-pBpF plasmid | KAN/CAM |
| Hs_BiPNBD_K123* | Lobe 1B | pet28a+ | BL21 DE3 +<br>pEVOL-pBpF plasmid | KAN/CAM |
| Hs_BiPNBD_K352* | Lobe 2A | pet28a+ | BL21 DE3 +<br>pEVOL-pBpF plasmid | KAN/CAM |
| Hs_BiPNBD_N380* | Lobe 2A | pet28a+ | BL21 DE3 +<br>pEVOL-pBpF plasmid | KAN/CAM |
| Hs_BiPNBD_R386* | Lobe 2A | pet28a+ | BL21 DE3 +<br>pEVOL-pBpF plasmid | KAN/CAM |
| Hs_BiPNBD_H252* | Lobe 2A | pet28a+ | BL21 DE3 +<br>pEVOL-pBpF plasmid | KAN/CAM |
| Hs_BiPNBD_E328* | Lobe 2B | pet28a+ | BL21 DE3 +<br>pEVOL-pBpF plasmid | KAN/CAM |
| Hs_BiPNBD_K294* | Lobe 2B | pet28a+ | BL21 DE3 +<br>pEVOL-pBpF plasmid | KAN/CAM |
| Hs_BiPNBD_E320* | Lobe 2B | pet28a+ | BL21 DE3 +<br>pEVOL-pBpF plasmid | KAN/CAM |
| Hs_BiPNBD_K272* | Lobe 2B | pet28a+ | BL21 DE3 +<br>pEVOL-pBpF plasmid | KAN/CAM |
| Hs_BiPNBD_H303* | Lobe 2B | pet28a+ | BL21 DE3 +<br>pEVOL-pBpF plasmid | KAN/CAM |

### Peptide Arrays

Peptide arrays were purchased from the MIT Biopolymers Laboratory. The tiling arrays were composed of 18-amino-acid peptides that were tiled along the BPTI sequence, with a three-amino-acid shift between adjacent spots. Arrays were incubated in 100% methanol for 10 min, then in Binding Buffer (50 mM HEPES pH 7.2, 150 mM NaCl, 0.02% Tween-20, 2 mM DTT) for 10 min three times, then with 500 nM MBP-IRE1-cLD for 1 hr at RT. The arrays were washed for 10 min with Binding Buffer three times to remove unbound protein. The bound protein was transferred to a nitrocellulose membrane using a semi-dry transfer apparatus and detected with anti-MBP antiserum (NEB). Peptide arrays were quantified using Max Quant such that binding intensity in each spot was normalized to max signal intensity in the array.

### Fluorescence Anisotropy

All peptides were synthesized by Genscript at >95% purity with an N-terminal 5-carboxyfluorescein (5-FAM) modification (**Table 4**). FAM-labelled peptides were held at a constant concentration of 100 nM in HKC buffer (25 mM HEPES pH 7.2, 75 mM KCl, and 2 mM CaCl_2_) and incubated with increasing amounts of either IRE1 cLD variants or BiP for 30 min at 25°C in a 20 µL reaction volume. Binding affinities were measured by the change in fluorescence anisotropy resulting from the titrated proteins as detected by a CLARIOstar BMG Labtech plate reader with excitation at 485 nm and emission at 525 nm. All assays were conducted in black, flat-bottomed, 384-well (low volume) microplates. Curves were fitted using GraphPad Prism software and the following equation: F_bound_ = r_free_ + (r_max_-r_free_)/(1 + 10^(LogK^1/2^-X)) where F_bound_ is the fraction of peptide bound, r_max_ and r_free_ are the anisotropy values at maximum and minimum plateaus, respectively, and X is the concentration of titrant protein. As this equation does not consider the equilibria between IRE1-cLD dimers/oligomers as protein concentration increases, affinity values are referred to as K_1/2_ in this work.

**Table 4.**
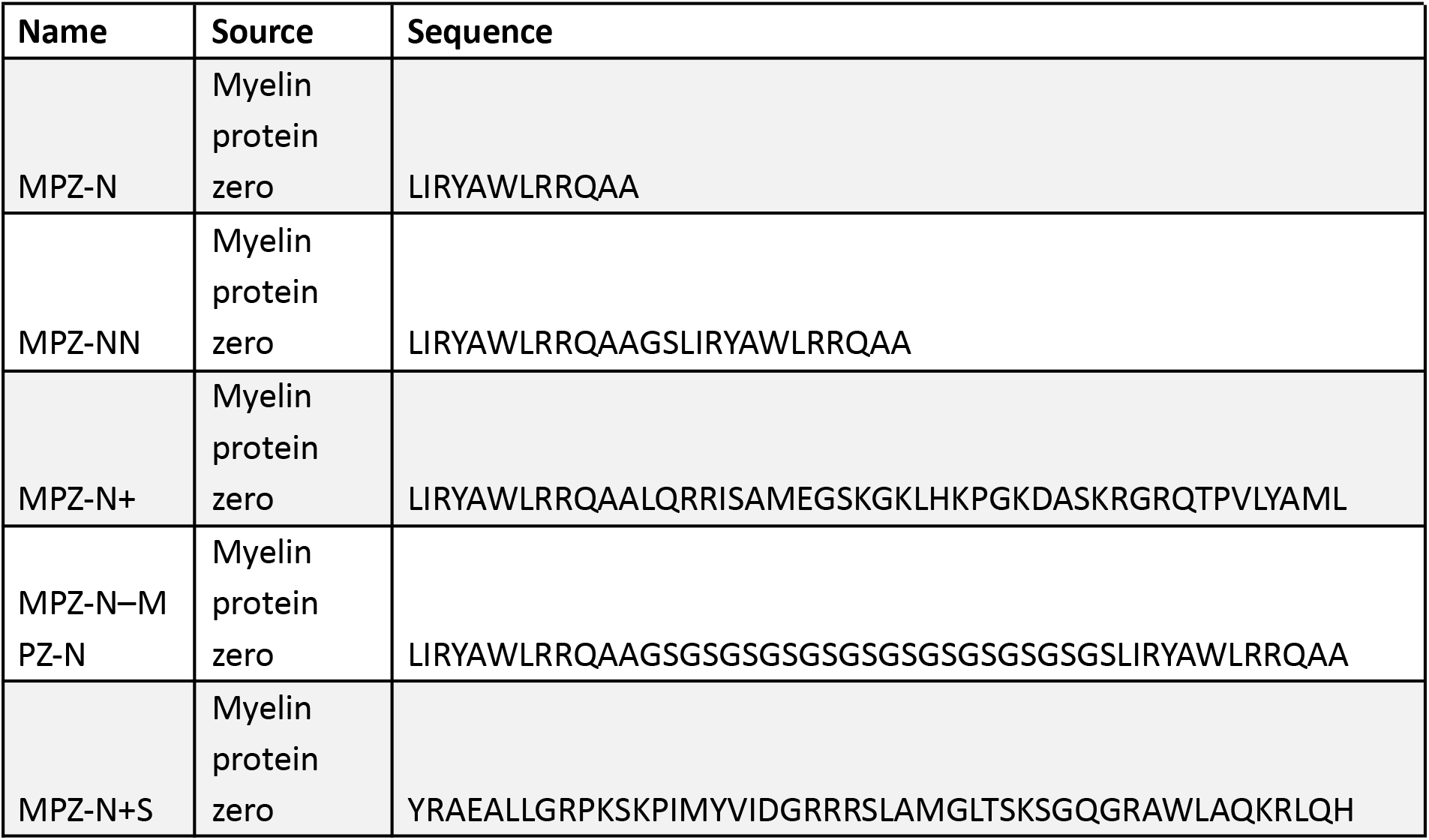
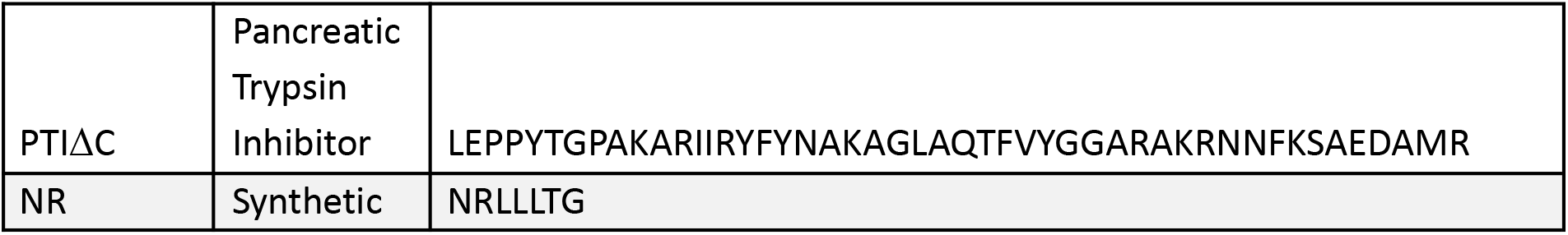
Summary of Peptide Sequences.

### OD350 Light Scattering Assay

The desired concentration of 5-FAM-labelled peptide or fully folded pancreatic trypsin inhibitor was added to 5 µM IRE1 cLD (or its variants) in HKC Buffer at 25°C in a 20 µL reaction volume. Oligomerization of IRE1 cLD was measured by the change in OD350 as detected by a SpectraMax5 plate reader measuring every 30 sec for 1 hr immediately upon addition of unfolded peptides. For filament assembly experiments with BiP NBD, IRE1 cLD was pre-incubated with the desired concentration of BiP NBD for 30 min at 25°C prior to the addition of unfolded peptide unless otherwise specified. All assays were conducted in clear, flat-bottomed, 384-well microplates.

### Negative Stain Microscopy

To assemble IRE1 cLD filaments, 5 µM WT cLD or its variants was incubated with 20 µM peptide for 1 hr at 25°C. Filament sample was applied at a volume of 2.5 µL to glow-discharged (30 sec, 30 mA) carbon-coated grids and incubated for 1 min. The sample was flushed away with 30 µL deionized water and followed by 30 µL uranyl formate (UF) stain. About 3 µL of UF stain was allowed to incubate for 30 sec before being flushed by another 30 µL of stain. This method of stain application was repeated two more times before filter paper side-blotting and gently wafting through the air to dry. Grids were imaged using a 120kV FEI Tecnai T12 equipped with a LaB6 filament and a Gatan UltraScan 895 4k CCD. Micrographs were collected at a magnification of 42,000x with -1.04 µm defocus. Micrograph measurements were taken using IMOD’s 3dmod program (79).

### Native Mass Spectrometry

Purified WT IRE1-cLD and BiP-NBD were buffer exchanged into 200 mM ammonium acetate by being run twice through 7 kDa MWCO Spin Desalting columns (Thermo Fisher Scientific). Nano electrospray was achieved in positive-ion mode on an Exactive Plus EMR orbitrap mass spectrometer (Thermo Fisher Scientific) using gold-coated capillaries (Thermo Fisher Scientific) and minimal backing pressure. The instrument was operated at a resolution of 35,000 (200 m/z), and an HCD cell trapping pressure set to 7, in-source trapping set to 10 eV. Spectra were acquired in the m/z range from 1000 to 20,000 with averaging of 5 microscans, injection time of 100 ms, and AGC target set to 1x10^6^. The remaining parameters were optimized in a sample dependent manner within the following ranges: capillary voltage (1.3 to 1.6 kV), and capillary temperature (200 to 250°C).

Mass spectra were deconvolved and plotted using UniDec software (81) with the peak detection threshold set to 0.1. The percent of IRE1-cLD protomers present in each species detected was calculated by weighting each species’ intensity value according to the number of protomers found in each species (1 in a monomer, 2 in a dimer, etc.). The percent of IRE1-cLD protomers in each species was then derived by dividing the weighted species intensity by the sum of all weighted species present.

### Analytical Size Exclusion Chromatography

IRE1 cLD variants diluted to 5 µM were loaded onto a Superdex 200 3.2/300 column (Cytiva) on a GE Healthcare Ettan liquid chromatography system pre-equilibrated with HKC Buffer (25 mM HEPES, 75 mM KCl, and 2 mM CaCl_2_). Samples were injected at a 50 µL volume and eluted over a 1.1 column volume isocratic elution using a 0.04 mL/min flow rate.

### IRE1 cLD Filament Centrifugation Assay

IRE1 cLD was pre-incubated at 5 µM with the desired concentration of 5-FAM labelled peptide for 30 min at 25°C in HKC Buffer before 40 µM of BiP NBD was added and allowed to incubate for a further 30 min at 25°C. Samples were then centrifuged at 1500xg for 1 min to separate the filamentous pellet from the supernatant. The pellet was washed 3X by sequentially resuspending in HKC Buffer and centrifuging at 1500xg for 1 min. Gel samples were taken for analysis by SDS-PAGE and Western Blotting.

### Western Blotting

All blots were performed on purified proteins incubated together and either cross-linked or not. Gel samples were made in 1X Novex Bolt LDS Sample Buffer supplemented with 5% Beta-mercaptoethanol and heated at 90°C for 10 min. Equal sample volumes were run on Invitrogen Bolt 4-12% Bis Tris Plus gels at 170 V for 45 min and transferred to PVDF membranes. Initial blocking was done by incubating for 2 hr at 4°C in 5% milk in PBST (1X Phosphate buffered saline and 0.1% TWEEN-20). Primary antibody staining was performed by incubating at the desired antibody dilution (**Table 5**) overnight at 4°C. The membranes were washed 1X for 5 min with 3% BSA in PBST and then a further 3X for 5 min in just PBST. Fluorescent secondary antibody staining was performed for 2 hr at 4°C in the dark before washing the membrane as described. Membranes were imaged on a Bio-Rad Chemidoc MP system using either the IRDye680RD or IRDye800CW channels.

**Table 5.** Summary of Antibodies for western blots.

| Antibody Target | Host | Dilution | Manufacturer |
| --- | --- | --- | --- |
| IRE1 alpha | Mouse | 1:10,000 | Santa Cruz Biotechnology |
| GRP78/BiP | Rabbit | 1:10,000 | Cell Signalling |
| Anti-mouse (fluorescent secondary) | Goat | 1:20,000 | LICOR Bio |
| Anti-rabbit (fluorescent secondary) | Goat | 1:20,000 | LICOR Bio |

### IRE1 cLD Dimer FRET Assay

AlexaFluor-546 Donor and AlexaFluor-647 Acceptor-IRE1 cLD (or its variants) were mixed at a 1:2 ratio (150 nM:300 nM) in HKC Buffer and incubated at 25°C in the dark for 30 min. Increasing amounts of unlabelled IRE1 cLD (or BiP) variants were then added and incubated for an hour at 25°C in the dark. Intactness of the cLD dimer interface was measured by the change in ‘FRET signal’ which was defined as the ratio between AF-546 emission (580 nm) and AF-647 emission (670 nm) following excitation by AF-546 (553 nm) as recorded by a CLARIOstar BMG Labtech plate reader. All assays were conducted in black, flat-bottomed, 384-well (low volume) microplates with top readings. Curves were fitted using GraphPad Prism software with the following equation: FRET_inhibition_ = FRET_min_ + (FRET_max_-FRET_min_)/(1 + 10^((LogIC50-X)*n_H_)) where FRET_inhibition_ is the inhibition of FRET signal across IF1, FRET_min_ and FRET_max_ are the FRET signal values at maximum and minimum plateaus, respectively, X is the concentration of titrant protein and n_H_ is the hill slope.

### Cross-linking and Mass Spectrometry-based Proteomics

#### Protein crosslinking and digestion

BiP-NBD and IRE1-cLD recombinant proteins were incubated at a 1:1 molar ratio (10 µM each) for 30 min at 25°C in a final volume of 40 mL of 25 mM HEPES, 150 mM KCl, and 2 mM MgCl_2_ (pH 7.2). tert-Butyl Disuccinimidyl Phenyl Phosphonate (TBDSPP or tBu-PhoX) was then added to a final concentration of 1 mM and the crosslinking reaction was carried out at 4°C for 1 hour before quenching with Tris pH 8.0 to a final concentration of 25 mM. Protein samples were reduced with 10 mM TCEP for 30 minutes at 25°C, 1500 rpm followed by alkylation with 40 mM 2-chloroacetamide for 30 minutes in the dark at 25°C, 1500 rpm. Proteins were digested with LysC and trypsin at a 1:40 ratio overnight at 37°C, 1000rpm.

#### Peptide C18 Clean-up and IMAC Enrichment

Digested peptides were acidified to 2% trifluoroacetic acid (TFA) and incubated for 1 hour at 25°C, 1500 rpm to deprotect the crosslinker prior to phospho-enrichment. Peptide samples were desalted by solid-phase extraction on an Agilent AssayMap Bravo using 5 μl C18 cartridges. Cartridges were activated with 50% acetonitrile (ACN), 0.1% TFA, and then equilibrated with 0.1% TFA in water. Samples were loaded, washed with 0.1% TFA, and eluted with 80% ACN, 0.1% TFA in water. Crosslinked peptides were enriched on the AssayMAP Bravo using 5 μL Fe(III)-NTA cartridges. The cartridges were primed with 0.1% TFA in ACN and equilibrated with 80% ACN, 0.1% TFA. Peptides were loaded, washed with 80% ACN/0.1% TFA, and eluted with 25 μL of 60% ACN, 25% NH_4_OH into a 96-well LoBind plate containing 25 μL of 10% formic acid in H_2_O to immediately neutralize the pH. The plate was immediately dried by centrifugation under vacuum and stored at -80°C. For LC-MS/MS analysis, the peptides were resuspended in 3% ACN:0.1% formic acid.

#### Mass Spectrometry Acquisition

Samples were measured on an Orbitrap Ascend mass spectrometer coupled to a Vanquish Neo HPLC with mobile phase A (0.1% formic acid in water) and B (80% ACN, 0.1% TFA in water). Peptides were separated on a 2-step linear gradient from 3-40% B over 90 mins followed by 40-60% B over 10 mins, at a flow rate of 400 nL/min on a 3rd Generation 25 cm Aurora Ultimate Columns (Ionopticks). A spray voltage of 1600V was applied and inlet capillary was heated to 275 C. 400,000 charges were accumulated in a maximum of 123 ms and measured in the Orbitrap at resolution of 60,000 from 380-1400 m/z. In a fixed cycle time of 3 seconds, peptides meeting the following criteria were selected for fragmentation, charge state from 3-8, minimum intensity of 50,000. Masses +-10 ppm were excluded from fragmentation for 60 seconds. Automatic gain control was used to select precursors at 200% of the normalized target, isolated using the quadrupole with a 1.6 m/z window, and fragmented using HCD with 25% and 30% normalized collision energies. Fragments were analyzed in the Orbitrap at a resolution of 30,000x with automatically determined scan range.

#### Database Search

Spectra files were searched using MSFragger (version 3.8) against the *E. coli* proteome (downloaded on June 23, 2023, UP000002032) including the relevant fusion protein sequences and a list of common contaminant proteins. Cysteine carbamidomethylation (57.02146) was set as fixed modification. Methionine oxidation (15.9949), protein N-term acetylation (42.0106), phosphorylation (79.96633 on S, T, Y), and PhoX crosslinker modifications (227.98238 and 331.0457 on K) were set as dynamic modifications. The proteins identified by MSFragger were combined into a FASTA file to generate a searchable library that was subsequently used with the XlinkX node 3.0 of the Proteome Discoverer 3.0 software (Thermo Fisher Scientific) to identify crosslinked peptides using the noncleavable open search algorithm. The XlinkX search was performed using cysteine carbamidomethylation as a fixed modification and methionine oxidation as a variable modification. PhoX (+209.97181 Da) was set as the crosslinker modification on lysine while also allowing protein N-terminus linkage with the following parameters: minimum peptide length 5, missed cleavages 2, precursor mass tolerance 10 ppm, fragment mass tolerance 20 ppm, and 5% FDR on CSM and cross-link levels. Crosslinks with threshold scores >40 and incidence >1 out of 3 independent sample replicates were visualized using ChimeraX (82).

### Filament Disassembly by OD350 Light Scattering

To preassemble cLD filaments, the desired concentration of 5-FAM-labelled peptide was added to 5 µM IRE1 cLD in HKC Buffer further supplemented with 10 mM MgCl_2_ and incubated for 1 hr at 25°C. BiP and ERDJ co-chaperone variants were then added before the mixture was pipetted out into clear, flat-bottomed, 384-well microplates with a reaction volume of 20 µL. Disassembly was initiated by spiking in 10 mM ATP into each well and tracked by the change in OD350 as detected by a SpectraMax5 plate reader measuring every 30 sec for 1 hr at 37°C. For experiments with an alternate BiP substrate, BiP was first incubated with NR peptide separately before addition to the preformed cLD filaments. Curve fittings to one phase exponential decay function and statistics were performed using GraphPad Prism software.

